# Disturbed engram network caused by NPTXs downregulation underlies aging-related memory deficits

**DOI:** 10.1101/2025.05.19.654996

**Authors:** Tao Jin, Yang Yang, Yu Guo, Yi Zhang, Qiumin Le, Nan Huang, Xing Liu, Jintai Yu, Lan Ma, Feifei Wang

## Abstract

Engram cells storing specific memories are allocated to separate neuronal ensembles, which preferentially recruit either excitatory or inhibitory inputs to drive precise memory expression. However, how these formed neuronal ensembles maintain their stability, and whether the disturbed stability contributes to aging-related memory deficits remain elusive. Here, we show that neuronal pentraxin1 (NPTX1) facilitates Kv7.2-mediated inhibition of *Fos^+^*ensemble hyperexcitability, thereby restricting its response to excitatory inputs from medial entorhinal cortex (MEC) and promoting memory expression in the fear context. Meanwhile, NPTX2 facilitates the perisomatic inhibition of the *Npas4^+^* ensemble by parvalbumin^+^ (PV^+^) interneurons, thus preventing fear memory overgeneralization. Pharmacological activation of Kv7.2 or chemogenetic activation of PV^+^ interneurons repaired memory deficits caused by engram specific NPTXs depletion. Contextual fear memory precision and NPTXs expression in dentate gyrus (DG) engram cells are decreased in aged mice. Overexpression of NPTX1 in *Fos^+^*ensemble or AMPAR binding domain of NPTX2 in *Npas4^+^* ensemble rescued memory imprecision. These findings elucidate that the coordination of NPTXs prevents engram ensembles from becoming hyperactive and provide a causal link between engram network destabilization and aging-associated memory deficits.

## Introduction

Memories are believed to be encoded in sparse neuronal ensembles (engram cells) that are activated by specific learning experiences^1,2^. Neurons recruited to engrams are more stably connected than those that are not^3^. Memory accessibility and precision are shaped by the plasticity of engram network, and require the long-lasting stabilization of the recruited engram network. However, the molecular mechanisms underlying the maintenance of the dormant, but stable engram network remain unknown. In addition, memory precision progressively declines with age, and is considered as one of the predominant hallmarks of aging-associated cognitive dysfunctions^4,5^, while it remains to be elucidated how the disturbed engram network leads to aging-related memory deficits.

Cell-adhesion molecules (CAMs) are a super family which play an irreplaceable role in synaptic plasticity through mediating the bidirectional organization of synaptic compartments^6^. NPTXs are a subfamily of CAMs primarily expressed in excitatory neurons. NPTXs consist of NPTX1, NPTX2 and their receptor NPTXR^7^. Pre-synaptic secreted NPTX1 and NPTX2 form disulfide-linked heteromultimers with post-synaptic NPTXR to promote the clustering of α-amino-3-hydroxy-5-methyl-4-isoxazolepropionic acid (AMPA) type glutamate receptors (AMPARs)^8,9^. Further studies identify NPTX1 functions as a pro-apoptotic protein^10^ that restricts excitatory synaptic transmission^11^. In contrast, NPTX2 maintains excitatory homeostasis by adaptively enhancing circuit inhibition^12,13^. These findings raise the possibility that NPTX1 and NPTX2 function as the potential molecular substrates for stabilizing the excitatory and inhibitory inputs onto distinct neuronal ensembles. NPTXs are also considered as prognostic biomarkers for neurodegeneration whose expressions are decreased in Alzheimer’s disease (AD), schizophrenia or even mild cognitive impairment (MCI) patients^14–16^. However, the causal link between NPTXs reduction-induced engram network destabilization and aging-related memory deficits remain unclear.

The tracing of neuronal circuits involved in specific memory formation is enabled by measuring the expression of selectively activated immediate early genes (IEGs). The DG engram cells within the equivalent memory contain functionally divergent *Fos^+^*and *Npas4^+^* neuronal ensembles, which drive precise memory expression by recruiting excitatory and inhibitory (E/I) circuits^17^. The expression of NPTX1 and NPTX2 are both abundant in DG^18^. In this study, we found that the maintenance of engram network stability requires NPTX1 and NPTX2 to coordinately stabilize the excitatory inputs to the *Fos^+^* ensemble and the inhibitory inputs to the *Npas4^+^* ensemble. Engram network hyperactivity, caused by NPTXs downregulation in DG engram ensembles contributed to aging-related contextual fear memory imprecision, including impaired memory expression in the fear context and overgeneralization in the non-fear context, which was rescued by restoring NPTXs expression in different engram ensembles. Our findings elucidate the critical role of NPTXs in stabilizing the formed engram network by controlling DG excitatory and inhibitory inputs, and establish a causal link between engram network destabilization and aging-associated memory deficits.

## Results

### NPTX1 restricts MEC excitatory inputs onto DG *F*-RAM ensemble

*Fos* and *Npas4-*dependent robust activity marking (*F*-RAM and *N*-RAM) reporter systems were used to label engram ensembles with mKate2 reporter by off Dox^17^. *Npas4-CreER^T^*^2^ mice in which the endogenous *Npas4* promoter drives the expression of *CreER^T^*^2^ were generated to label *Npas4*^+^ ensembles with EGFP reporter by 4-hydroxytamoxifen (4-OHT) intraperitoneal (i.p.) injection (Figure. S1A-1D). A combination of these two activity-dependent labelling strategies (Figure. S1E) was used simultaneously to label engrams activated by contextual fear conditioning (CFC, 0.5 mA, 1 s, 3 trials). The EGFP^+^ neurons showed substantial overlap with mKate2^+^ *N*-RAM ensemble while showed minimal colocalization with mKate2^+^ *F*-RAM ensemble (Figure. S1F-1G). These results confirm that *Fos^+^* and *Npas4^+^*engram cells in DG labeled by *F*-RAM and *N*-RAM reporters are distinct populations.

The upstream inputs of *F*– and *N*-RAM neuronal ensembles from medial septum (MS), horizontal limb of the diagonal band (HDB), perirhinal cortex (PRh), lateral entorhinal cortex (LEC), medial entorhinal cortex (MEC) and DG were detected by rabies virus (RV) tracing (Figure 1A-1B). *F*-RAM ensemble formed more connections with MS and MEC, while *N*-RAM ensemble received more inputs from DG local neurons (Figure 1C). The majority of dsRed^+^ neurons in MS were co-labelled with the cholinergic neuron marker choline acetyltransferase (ChAT, Figure 1D-1E), and DG *F*-RAM neurons received more excitatory inputs from MEC than *N*-RAM neurons (Figure 1F-1G).

**Figure 1.**
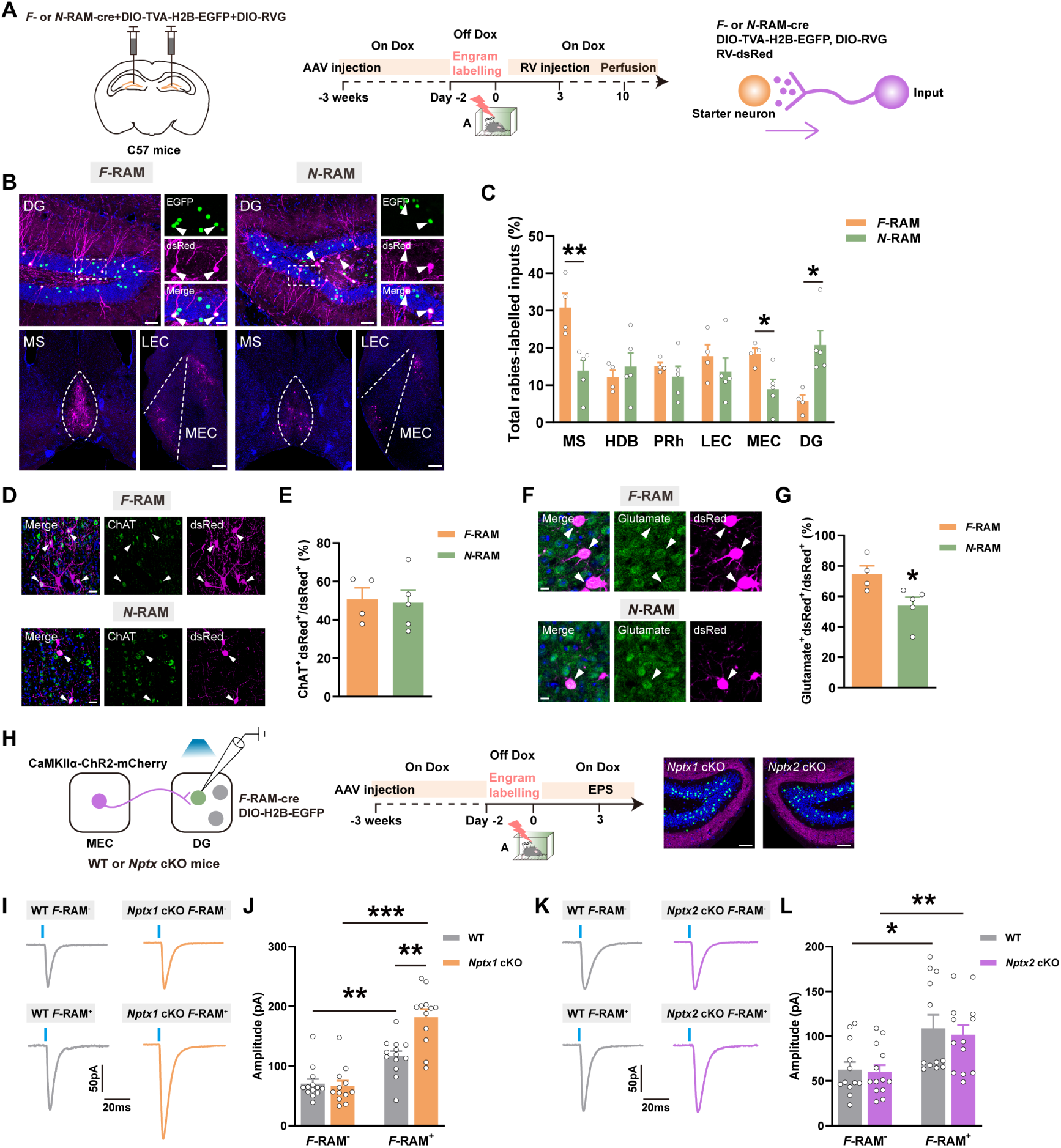
Rabies virus tracing of the upstream inputs to DG *F*– and *N*-RAM neuronal ensembles and the effects of *Nptxs* depletion on response to MEC excitatory inputs. (A) Diagram of virus injection and experimental scheme to trace upstream inputs of DG *F*-RAM and *N*-RAM ensembles. (B) Representative confocal images of DG starter neurons in *F*-RAM, *N*-RAM ensembles and their respective upstream inputs. Green: *F*-RAM or *N*-RAM engram cells, EGFP, Purple: dsRed, Blue: DAPI. White arrows indicate the starter neurons and DG local inputs. Scale bar: top left, 50 μm, top right, 10 μm, bottom, 100 μm. (C) The percentages of total inputs from each upstream site relative to total quantified inputs. (D) Representative confocal images of upstream MS dsRed^+^ cells immunostaining with ChAT. Green: ChAT, Purple: dsRed, Blue: DAPI. White arrows indicate the colocalized cells. Scale bar: 10 μm. (E) The percentages of colocalized cells in MS total dsRed^+^ cells. (F) Representative confocal images of upstream MEC dsRed^+^ cells immunostaining with glutamate. Green: glutamate, Purple: dsRed, Blue: DAPI. White arrows indicate the colocalized cells. Scale bar: 5 μm. (G) The percentages of colocalized cells in MEC total dsRed^+^ cells. (H) Diagram of AAV injection, experimental scheme and representative expression of ChR2 in MEC projection neurons and DG engram cells. Green: *F*-RAM engram cells of *Nptx1* and *Nptx2* cKO mice, EGFP, Purple: MEC projections, mCherry, Blue: DAPI. Scale bar: 50 μm. (I, K) Representative traces of oEPSC recorded from WT and *Nptxs* cKO mice. (J) The quantification of oEPSC amplitudes recorded from WT and *Nptx1* cKO mice. (L) The quantification of oEPSC amplitudes recorded from WT and *Nptx2* cKO mice. Data are presented as mean ± S.E.M; \**P* < 0.05, \*\**P* < 0.01, \*\*\**P* < 0.001.

NPTXs were found to be predominantly involved in E/I synaptic homeostasis rather than cholinergic signaling^9,19^. To test whether NPTXs modulate the stability of the synaptic transmission between MEC and DG *Fos^+^* ensemble, *Nptx1^fl/fl^*, *Nptx2 ^fl/fl^* conditional knockout (*Nptx1* cKO, *Nptx2* cKO) mice were generated (Figure S2A-2B). Ribosome-associated transcripts of *F*-RAM and *N*-RAM ensembles were enriched from *Nptxs* cKO mice and their WT littermates (Figure S2C-2D), and qPCR against *Nptx1* exon 3 and *Nptx2* exon 2 was performed to confirm the knockout efficiency in *F-* or *N-*RAM ensembles. *Nptx2* expression was unaffected when *Nptx1* was depleted and vice versa (Figure S2E-2L). Immunostaining of NPTX1 and NPTX2 further confirmed the knockout efficiency in DG engram ensembles (Figure S3).

*AAV-CaMKIIα-ChR2-mCherry* was then injected into MEC of *Nptx1* cKO, *Nptx2* cKO mice and their WT littermates, and the virus mixture of *AAV-F-RAM-Cre* with Cre recombinase-dependent *AAV-DIO-H2B-EGFP* was injected into DG to knock out *Nptxs* in *Fos^+^* ensemble. Opto-evoked excitatory post-synaptic currents (oEPSCs) from MEC were recorded on EGFP^-^ (*F*-RAM^-^) and EGFP^+^ (*F*-RAM^+^) cells (Figure 1H). The amplitude of oEPSCs on EGFP^+^ cells was larger in the *Nptx1* cKO group (Figure 1I-1J), whereas this effect was not found in the *Nptx2* cKO mice (Figure 1K-1L). These data indicate that NPTX1 plays an important role in dampening the excitatory synaptic transmission between MEC and DG *Fos^+^*ensemble.

### NPTX1 facilitates *I*_M_ current and membrane expression of Kv7.2 in DG *F*-RAM ensemble

Action potential (AP) recordings were performed to examine whether neuronal excitability of the engram ensembles was changed after *Nptxs* knockout (Figure 2A-2C). *F*-RAM engram cells lacking *Nptx1,* but not *Nptx2,* exhibited upward shift of the neuronal spiking curve after step-increment current injections, lower resting membrane potential (RMP) and rheobase (Figure 2D-2K), indicating increased excitability of *F*-RAM engram cells after *Nptx1* knockout. KCNQ2 (Kv7.2) pairs with the KCNQ3 (Kv7.3) subunit to form KCNQ2/3 heterotetramers, which are the primary molecular substrate of M-current (*I*_M_) involved in regulating neuronal excitability^20,21^. Application of the KCNQ2/3-selective blocker XE991 resulted in a reduction of *I*_M_ amplitude on DG *F*-RAM engram cells (Figure 2L-2M). *Nptx1* depletion exerted the similar inhibitory effect of XE991 on *I*_M_ amplitude (Figure 2N-2O). Co-immunoprecipitation (Co-IP) of the DG lysate confirmed the interaction between NPTX1 and Kv7.2 (Figure 2P). Immunostaining analysis further revealed that *Nptx1* ablation decreased Kv7.2 membrane expression in *F*-RAM engram cells (Figure 2Q-2R), which was not observed in *Nptx2* cKO group (Figure 2S-2T). Taken together, these results indicate that NPTX1 restrains the excitatory synaptic transmission between MEC and DG *Fos^+^* ensemble by facilitating Kv7.2 membrane expression-mediated inhibition of neuronal hyperexcitability.

**Figure 2.**
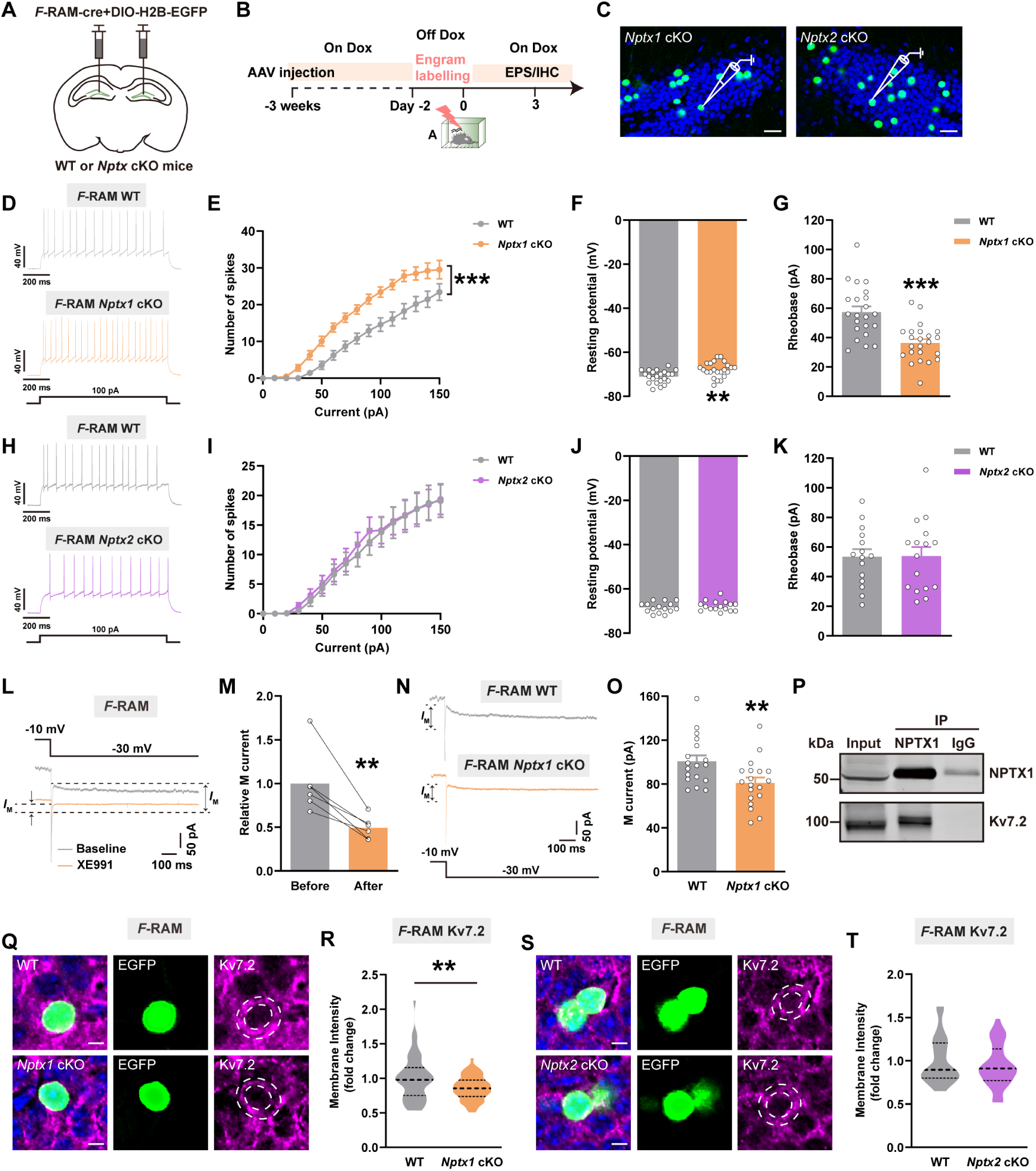
The effects of *Nptx1* or *Nptx2* depletion on engram excitability and membrane expression of Kv7.2. (A) Diagram of AAV injection. (B) Experimental scheme to label *F*-RAM, *N*-RAM engram ensembles. EPS: Electrophysiology, IHC: Immunohistochemistry. (C) Representative expression of *F*-RAM engram cells in DG of *Nptx1* and *Nptx2* cKO mice. Green: *F*-RAM engram cells of *Nptx1* and *Nptx2* cKO mice, EGFP, Blue: DAPI. Scale bar: 20 μm. (D, H) Representative AP traces induced by depolarizing current injections (100 pA) recorded from WT and *Nptxs* cKO mice. (E) The input-output curves of AP spikes versus injected currents of WT and *Nptx1* cKO mice. (F) The RMP of WT and *Nptx1* cKO mice. (G) The rheobase of WT and *Nptx1* cKO mice. (I) The input-output curves of AP spikes versus injected currents of WT and *Nptx2* cKO mice. (J) The RMP of WT and *Nptx2* cKO mice. (K) The rheobase of WT and *Nptx2* cKO mice. (L) Representative traces of *I*_M_ current recorded before and after application of XE991. (M) The normalized *I*_M_ current recorded at –30 mV before and after application of XE991. (N) Representative traces of *I*_M_ current recorded from WT and *Nptx1* cKO mice. (O) The average *I*_M_ current amplitudes recorded at –30 mV from WT and *Nptx1* cKO mice. (P) Immunoblotting of NPTX1 co-immunoprecipitates with Kv7.2 in DG. (Q, S) Representative confocal images of engram cells colocalizing with Kv7.2. Green: *F*-RAM engram cells, EGFP, Purple: Kv7.2, Blue: DAPI. Dashed white lines outline cell membrane. Scale bar: 5 μm. (R) The quantification of membrane Kv7.2 fluorescence intensity of WT and *Nptx1* cKO neurons. (T) The quantification of Kv7.2 fluorescence intensity of WT and *Nptx2* cKO neurons. Data are presented as mean ± S.E.M; \*\**P* < 0.01, \*\*\**P* < 0.001.

### NPTX2 in *N*-RAM ensemble facilitates the perisomatic inhibition by PV^+^ interneurons in DG

Rabies tracing confirmed *N*-RAM ensemble received more inputs from DG local neurons (Figure 1C), leading us to wonder whether NPTX2 was specifically involved in *Npas4^+^* engram-DG local neuron synaptic connection. Immunostaining showed that parvalbumin^+^ (PV^+^) and somatostatin^+^ (SST^+^) interneurons, two major subtypes of GABAergic interneurons in DG^22^ that separately target perisomatic compartments and distal apical dendrites of granule cells^23^ were both recruited by DG *Fos^+^* and *Npas4^+^* engram cells after CFC. However, *Npas4^+^* engram cells received more PV^+^ but not SST^+^ inputs in DG (Figure 3A-3B, Figure S4A-4B).

**Figure 3.**
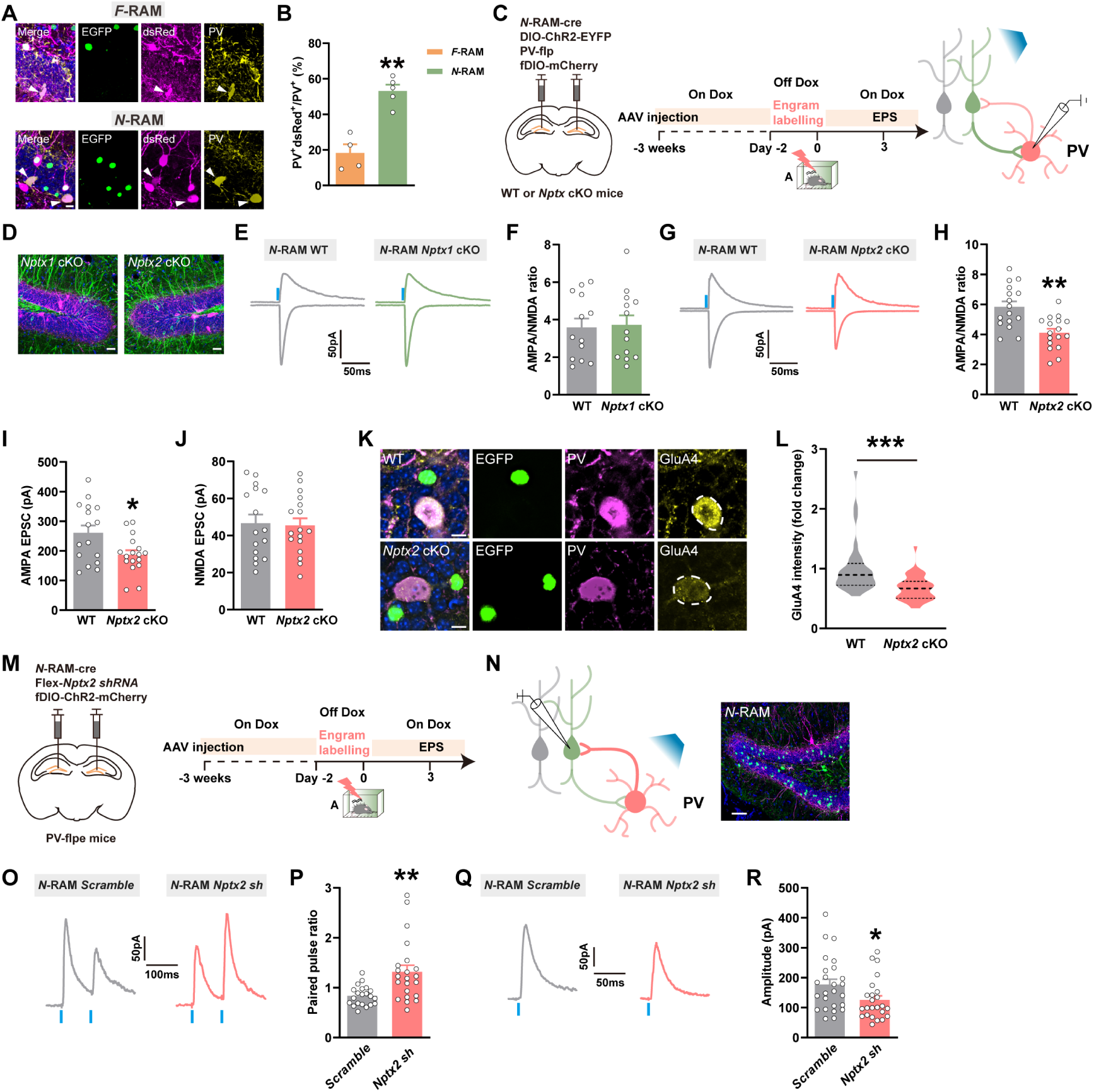
The effects of *Nptx2* depletion on the functional connection between DG local PV^+^ interneurons and *N*-RAM ensemble. (A) Representative confocal images of DG local dsRed^+^ cells immunostaining with PV. Green: EGFP, Purple: dsRed, Yellow: PV, Blue: DAPI. White arrows indicate the dsRed^+^ PV^+^ colocalized cells. Scale bar: 5 μm. (B) The percentages of colocalized cells in DG total PV^+^ cells. (C) Diagram of AAV injection and experimental scheme to label *N*-RAM engram ensembles. (D) Diagram of representative expression of engram cells and PV^+^ interneurons. Green: *N*-RAM engram cells of *Nptx1* and *Nptx2* cKO mice, EYFP, Purple: PV^+^ interneurons, mCherry, Blue: DAPI. Scale bar: 10 μm. (E, G) Representative traces of opto-evoked AMPA-EPSC and NMDA-EPSC recorded from WT and *Nptxs* cKO mice. (F) The average A/N ratio recorded from WT and *Nptx1* cKO mice. (H) The average A/N ratio recorded from WT and *Nptx2* cKO mice. (I, J) The average AMPA-EPSC and NMDA-EPSC amplitudes recorded from WT and *Nptx2* cKO mice. (K) Representative confocal images of GluA4 colocalizing with PV^+^ interneurons. Green: *N*-RAM engram cells, Purple: PV^+^ interneurons, Yellow: GluA4. Scale bar: 10 μm. (L) The quantification of GluA4 membrane expression on PV^+^ interneurons in WT and *Nptx2* cKO mice. (M) Diagram of AAV injection and experimental scheme to label *N*-RAM engram ensemble. (N) Diagram of photostimulation and whole-cell patch clamp recordings (left) and representative expression of *N*-RAM engram cells and PV^+^ interneurons (right). Green: *N*-RAM engram cells, EGFP, Purple: PV^+^ interneurons, mCherry, Blue: DAPI. Scale bar: 20 μm. (O, Q) Representative traces of opto-evoked PPR and oIPSC recorded from *Scramble* and *Nptx2 shRNA* neurons. (P, R) The quantification of PPR and oIPSC amplitudes recorded from *Scramble* and *Nptx2 shRNA* neurons. Data are presented as mean ± S.E.M; \**P* < 0.05, \*\**P* < 0.01, \*\*\**P* < 0.001.

To test the involvement of NPTX2 in the synaptic transmission between *N*-RAM ensemble and DG interneurons, the viral mixture of *AAV-N-RAM-Cre, AAV-DIO-ChR2-EYFP, AAV-PV/SST-Flp* and Flp recombinase-dependent *AAV-fDIO-mCherry* was injected into DG of *Nptx1* cKO, *Nptx2* cKO mice and WT littermates. The specificity of *AAV-SST-Flp* and *AAV-PV-Flp* was confirmed by immunostaining with anti-SST and anti-PV antibodies (Figure S4C-4D, Figure S5A). Opto-evoked paired-pulse ratio (PPR) and AMPAR/N-methyl-D-aspartate receptor (NMDAR) (A/N) ratio were recorded on SST^+^ or PV^+^ interneurons according to mCherry expression, morphology and location (Figure 3C-3D, Figure S4E-4F). *Nptxs* depletion in *N*-RAM ensemble reduced pre-synaptic glutamate release (Figure S4G-4H, S4K-4L, Figure S5B-5E,), consistent with previous studies that NPTXs family promotes pre-synaptic glutamate release^19,24^. *Nptx2* knockout in *N*-RAM ensemble exerted no effect on A/N ratio of SST^+^ interneurons (Figure S4I-4J, S4M-4N), whereas *Nptx2,* but not *Nptx1* depletion in *N*-RAM ensemble significantly decreased AMPAR-mediated EPSC and A/N ratio on PV^+^ interneurons (Figure 3E-3J). Previous studies found that NPTX2 binds AMPAR subunit GluA4 on PV^+^ interneurons to regulate network excitatory/inhibitory balance^12,25^. Immunostaining confirmed that *Nptx2* depletion in *N*-RAM ensemble decreased the expression of GluA4 on PV^+^ interneurons (Figure 3K-3L).

To examine whether NPTX2-dependent plasticity applies to GABAergic CCK^+^ cells, the viral mixture of *AAV-N-RAM-Cre, AAV-DIO-ChR2-EYFP* with *AAV-vGAT2-Flp*-dependent expression of *AAV-CCK-fDIO-mCherry* was injected into DG of WT and *Nptx2* cKO mice to label the GABAergic CCK^+^ cells (Figure S6A-6C). Opto-evoked PPR and A/N ratio were recorded on mCherry^+^ interneurons, and no differences were detected after *Nptx2* depletion in *N*-RAM ensemble (Figure S6D-6G). These data suggest that NPTX2-dependent interneuron plasticity was specific to PV^+^ interneurons. The disturbed PV^+^ interneuron plasticity caused by *Nptx2* depletion led us to speculate whether the inhibitory inputs received by *N*-RAM ensemble was also affected. A viral mixture of *AAV-fDIO-ChR2-mCherry*, *AAV-N-RAM-Cre* and *AAV-Flex-EGFP-Nptx2 shRNA (Nptx2 sh)* was injected into DG of *PV-Flpe* mice, and opto-evoked inhibitory post-synaptic currents (oIPSCs) and PPR were recorded on *N*-RAM cells (Figure 3M-3N). The increased PPR in the *Nptx2 shRNA* group indicates the reduction of pre-synaptic γ-aminobutyric acid (GABA) release from PV^+^ interneurons (Figure 3O-3P), leading to the reduced inhibition *Npas4^+^* engram cell received (Figure 3Q-3R). Taken together, these data indicate that NPTX2 facilitates the inhibitory synaptic transmission between DG *Npas4^+^* engram cells and PV^+^ interneurons by the AMPAR-dependent mechanism.

### NPTX1 in DG *F*os^+^ ensemble and NPTX2 in DG *Npas4^+^*ensemble facilitate the precise expression of contextual fear memory

To investigate the involvement of NPTXs-dependent maintenance of engram network stability on behavioral outputs, the viral mixture of *AAV-F-RAM-Cre* or *AAV-N-RAM-Cre* combined with *AAV-DIO-EYFP* was injected into DG of young *Nptx1* cKO, *Nptx2* cKO mice and their respective WT littermates. Mice were tested on day 3 in context A and context C after fear conditioning (Figure 4A-4C, 4H-4J). The freezing level during conditioning was not different between *Nptx1* cKO, *Nptx2* cKO mice and their WT littermates (Figure S7A-7D). Interestingly, *Nptx1* depletion in *F*-RAM ensemble decreased freezing in context A (Figure 4D-4E), whereas *Nptx2* depletion in *F*-RAM neuronal ensemble caused no differences in freezing level in both contexts (Figure 4F-4G). *Nptx1* knockout in DG *N*-RAM ensemble did not affect the freezing level in either context (Figure 4K-4L), whereas *Nptx2* knock out in DG *N*-RAM ensemble increased the freezing level in the non-fear context C on day 3, indicating overgeneralization of contextual fear memory (Figure 4M-4N).

**Figure 4.**
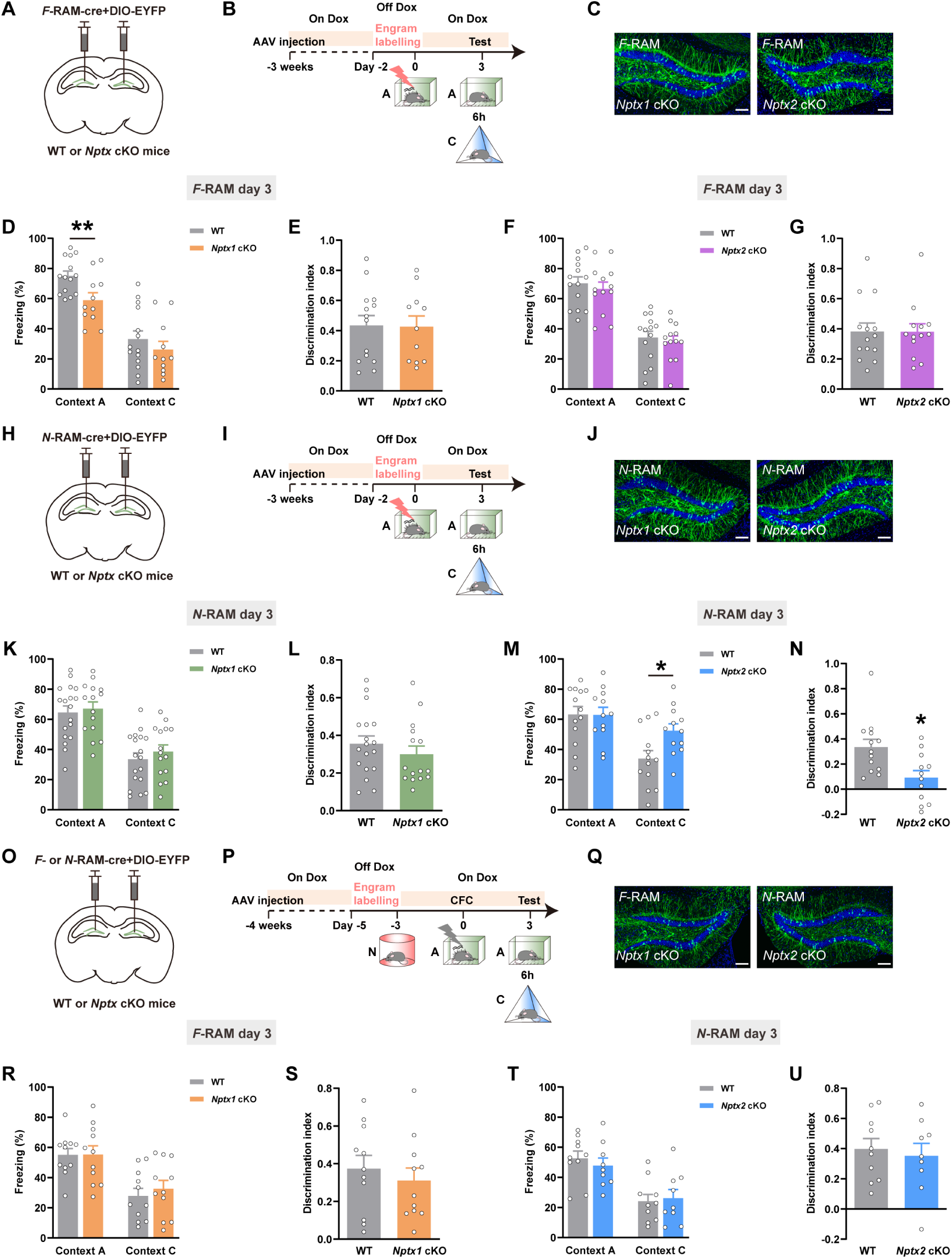
The effects of *Nptxs* depletion in *F*-RAM and *N*-RAM ensembles on the expression of contextual fear memory. (A, H, O) Diagram of AAV injection. (B, I, P) Experimental scheme of CFC and memory retrieval. (C, J, Q) Representative expression of *F*– and *N*-RAM engram cells in DG. Green: *F*-RAM or *N*-RAM ensemble, EYFP, Blue: DAPI. Scale bar: 100 μm. (D, E) The freezing percentage and discrimination index of WT and *Nptx1* cKO mice tested in context A and context C at day 3 (*F*-RAM). (F, G) The freezing percentage and discrimination index of WT and *Nptx2* cKO mice tested in context A and context C at day 3 (*F*-RAM). (K, L) The freezing percentage and discrimination index of WT and *Nptx1* cKO mice tested in context A and context C at day 3 (*N*-RAM). (M, N) The freezing percentage and discrimination index of WT and *Nptx2* cKO mice tested in context A and context C at day 3 (*N*-RAM). (R, S) The freezing percentage and discrimination index of WT and *Nptx1* cKO mice tested in context A and context C at day 3 (*F*-RAM). (T, U) The freezing percentage and discrimination index of WT and *Nptx2* cKO mice tested in context A and context C at day 3 (*N*-RAM). Data are presented as mean ± S.E.M; \**P* < 0.05, \*\**P* < 0.01.

In addition, when mice were exposed to a novel context (context N) quite different with context A or context C, *Nptx1* or *Nptx2* was knocked out in randomly labeled *F*-or *N*-RAM neurons (Figure 4O-4Q), and no differences in freezing were found between *Nptxs* cKO and WT controls (Figure S7E-7F, Figure 4R-4U), indicating the fear context-specific effect of *Nptxs*. Moreover, *Nptxs* depletion in both ensembles had no effect on mice’s locomotion, anxiety or depression levels (Figure S8-S10).

Taken together, these data suggest that the precise expression of contextual fear memory, including high freezing level in the fear context but low freezing level in the non-fear context requires the coordination of NPTX1 and NPTX2 functioning in distinct ensembles.

### Pharmacological activation of Kv7.2 or chemogenetic activation of PV^+^ interneurons ameliorates memory deficits induced by NPTXs depletion in DG engrams

The Kv7.2 activator retigabine, a clinically used anticonvulsant^26^, was i.p. injected 30 min before mice were tested in the fear context A and non-fear context C (Figure 5A-5C). Retigabine had no effect on the freezing level of WT mice in either context, whereas it restored the reduced freezing level in context A when NPTX1 was depleted in *F*-RAM ensemble but not in *N*-RAM ensemble (Figure 5D-5G, Figure S11A-11B). To rescue memory overgeneralization induced by *Nptx2* knockout in *N*-RAM ensemble, the designer receptors exclusively activated by designer drugs (DREADDs) were used to activate PV^+^ interneurons. The viral mixture of *AAV-N-RAM-Cre*, *AAV-Flex-EGFP-Nptx2 shRNA* and *AAV-fDIO-hM3Dq-mCherry* was injected into the DG of *PV-Flpe* mice and clozapine N-oxide (CNO) was administrated by i.p. injection 30 min before the mice were tested in context A and context C (Figure 5H-5J). Chemogenetic activation of local DG PV^+^ interneurons had no effect on the freezing level of control group infected with *Scramble shRNA*, while it reduced the freezing level of mice in context C when *Nptx2* was knocked down in *N*-RAM ensemble instead of *F*-RAM ensemble (Figure 5K-5N, Figure S11C-11D).

**Figure 5.**
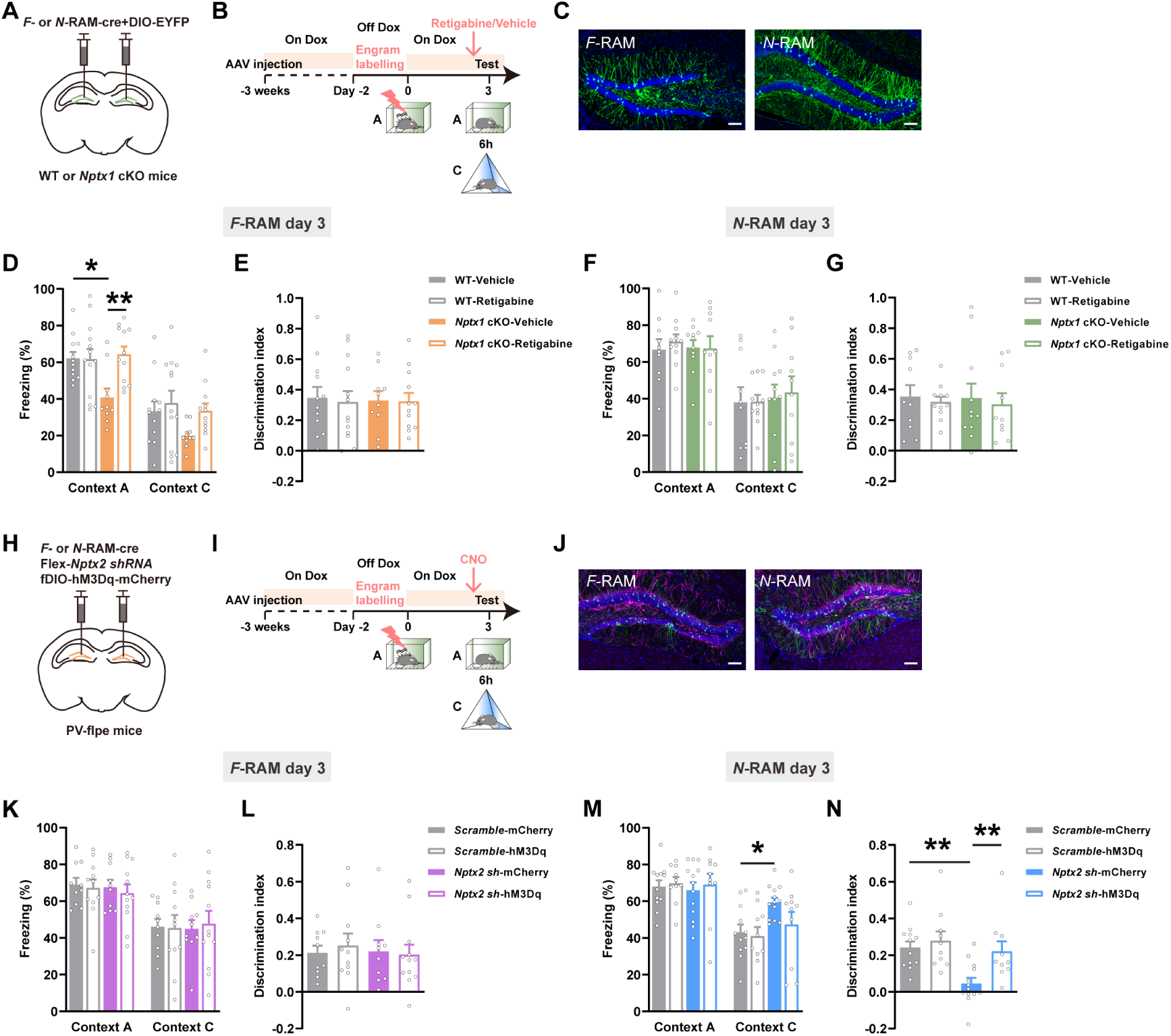
The effects of activating Kv7.2 or DG PV^+^ interneurons on memory deficits induced by *Nptxs* depletion in DG engram ensembles. (A, H) Diagram of AAV injection. (B, I) Experimental scheme of memory retrieval test. (C) Representative images of *F*– and *N*-RAM engram cells in DG. Green: *F*-RAM and *N*-RAM ensembles, EYFP, Blue: DAPI. Scale bar: 100 μm. (D, E) The freezing percentage and discrimination index of WT-Vehicle, WT-Retigabine, *Nptx1* cKO-Vehicle and *Nptx1* cKO-Retigabine mice (*F*-RAM). (F, G) The freezing percentage and discrimination index of WT-Vehicle, WT-Retigabine, *Nptx1* cKO-Vehicle and *Nptx1* cKO-Retigabine mice (*N*-RAM). (J) Representative images of *F*– and *N*-RAM engram cells in DG. Green: *F*-RAM and *N*-RAM ensembles, EGFP, Purple: PV^+^ interneurons, mCherry, Blue: DAPI. Scale bar: 100 μm. (K, L) The freezing percentage and discrimination index of *Scramble*-mCherry, *Scramble*-hM3Dq, *Nptx2 sh*-mCherry and *Nptx2 sh*-hM3Dq groups (*F*-RAM). (M, N) The freezing percentage and discrimination index of *Scramble*-mCherry, *Scramble*-hM3Dq, *Nptx2 sh*-mCherry and *Nptx2 sh*-hM3Dq groups (*N*-RAM). Data are presented as mean ± S.E.M; \**P* < 0.05, \*\**P* < 0.01.

### Downregulation of *Nptxs* in DG engrams in aged mice

Maladaptive changes in the expression of functional synaptic protein leading to neuronal network destabilization and cognitive impairment, are central hallmarks associated with physiological brain aging^27^. NPTXs are considered as biomarkers of synaptic dysfunction and cognitive impairment^7,16^. To investigate whether *Nptxs* expression in DG changes during physiological aging, young (3m) and aged (18m) mice were sacrificed 15 min, 30 min, 60 min and 120 min after CFC or under homecage (HC) condition, smFISH was used to assess their mRNA expression (Figure 6A-6D, S12A-12D).

**Figure 6.**
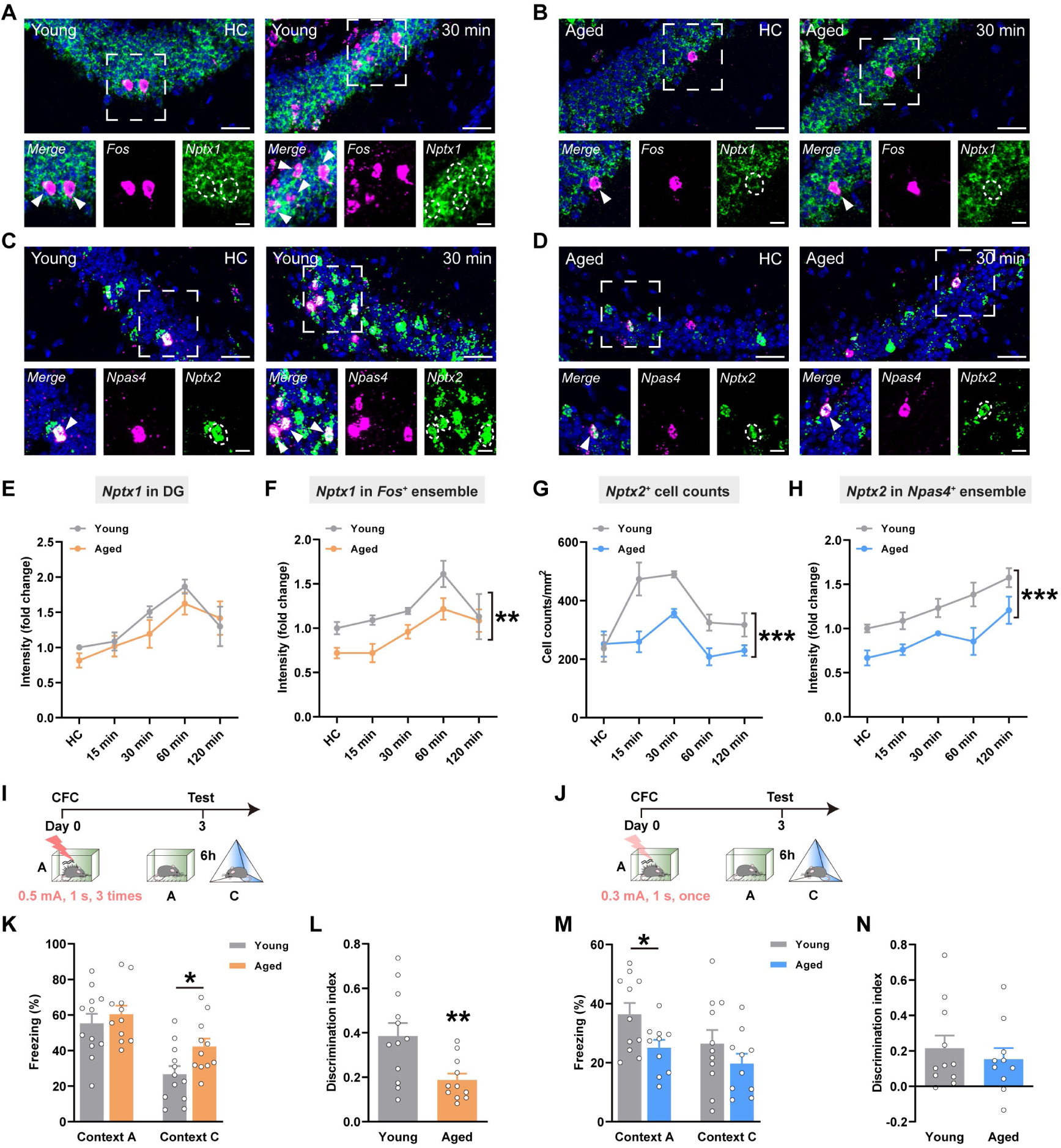
*Nptxs* expression in DG engram cells and contextual fear memory in young and aged mice. (A, B) Representative confocal images of *Nptx1* colocalizing with *Fos* under HC condition and 30 min after CFC in DG of young and aged mice. Green: *Nptx1*, Purple: *Fos*, Blue: DAPI. White arrows indicate the colocalized cells. Scale bar: top, 30 μm, bottom, 10 μm. (C, D) Representative confocal images of *Nptx2* colocalizing with *Npas4* under HC condition and 30 min after CFC in DG of young and aged mice. Green: *Nptx2*, Purple: *Npas4*, Blue: DAPI. White arrows indicate the colocalized cells. Scale bar: top, 30 μm, bottom, 10 μm. (E) The fluorescence intensity of overall *Nptx1* mRNA in DG. (F) The fluorescence intensity of *Nptx1* mRNA in *Fos^+^* ensemble at HC, 15 min, 30 min, 60 min and 120 min after CFC. (G) The *Nptx2^+^* cell counts in DG. (H) The fluorescence intensity of *Nptx2* mRNA in *Npas4^+^* ensemble at HC, 15 min, 30 min, 60 min and 120 min after CFC. (I, J) Experimental scheme of CFC to test memory expression. (K, M) The freezing percentages of young and aged mice. (L, N) The discrimination indexes of young and aged mice. Data are presented as mean ± S.E.M; \**P* < 0.05, \*\**P* < 0.01, \*\*\**P* < 0.001.

*Fos^+^* and *Npas4^+^* cell numbers after CFC were decreased in the DG of aged mice, especially at 30 and 60 min, indicating decreased activation of neurons during learning in aged mice (Fig S12E-12F). *Nptx1* was expressed in all DG granule cells, although the overall *Nptx1^+^* fluorescence intensity was not significantly decreased in aged mice (Figure 6E), the specific expression of *Nptx1* in *Fos^+^* and *Npas4^+^* engram cells were reduced (Figure 6F, Figure S12G). NPTX2 was also seen as a neuronal IEG protein^14^, both *Nptx2^+^*cell numbers and the average fluorescence intensity of *Nptx2 mRNA* in *Fos^+^* and *Npas4^+^* engram cells were decreased in aged mice (Figure 6G-6H, Figure S12H). These findings indicate the downregulation of *Nptx1* and *Nptx2* in *Fos^+^* and *Npas4^+^* engram cells in aged mice.

The expression of contextual fear memory in young and aged mice were further assessed. When exposed to strong fear conditioning (0.5 mA, 1 s, 3 trials), aged mice froze more in the non-fear context C, suggesting memory overgeneralization (Figure 6I, 6K-6L). However, when exposed to mild fear conditioning (0.3 mA, 1 s, 1 trial), aged mice froze less in the fear context A (Figure 6J, 6M-6N), suggesting memory retrieval impairment. These results demonstrate that aging impairs the precise expression of contextual fear memory.

### NPTXs overexpression in DG engrams rescued memory deficits in aged mice

Consistent with the decreased number of *Fos^+^* and *Npas4^+^* cells in DG after CFC, the number of cells labelled by the *F*– and *N*-RAM systems were both reduced in DG of aged mice (Figure S12I-12J). Ribosome-associated transcripts of *F*-RAM and *N*-RAM ensembles activated in the home cage or by CFC were analyzed (Figure S13A). Gene ontology (GO) analysis revealed clusters of differentially expressed genes (DEGs) related to synaptic plasticity and cell communication between *F*-RAM and *N*-RAM ensembles, which were more significant after CFC (Figure S13B). In addition, a minority of DEGs overlapped between *F*– and *N*-RAM ensembles during aging (Figure S13C-13D). DEGs in *F*-RAM during aging were preferentially enriched in pathways regulating neuronal excitability, including membrane potential and potassium ion import, whereas DEGs in *N*-RAM during aging were preferentially enriched in AMPAR-mediated glutamatergic transmission and others (Figure S13E), suggesting different pathways involved in engram network plasticity during aging between *F*– and *N*-RAM ensembles.

Decreased Kv7.2 membranes expression on c-Fos^+^ engram cells (Figure 7A-7B) and NPTX1-Kv7.2 interaction in DG (Figure 7C-7D, S14A) were observed in aged mice. AAV-DIO-Nptx1-EYFP was constructed to overexpress NPTX1 in a Cre-dependent way (Figure S14B). Aged mice were subjected to the mild fear conditioning protocol (0.3 mA, 1 s, one shock). Overexpression of NPTX1 in *F*-RAM ensemble, but not in *N*-RAM ensemble increased the freezing level of aged mice in the fear context A (Figure 7E-7K, Figure S14C-14D). Decreased GluA4 expression on PV^+^ interneurons and PV^+^ fluorescence intensity surrounding Npas4^+^ engram cells were observed in aged mice (Figure 7L-7O), which are consistent with the phenotype of *Nptx2* depletion in young mice (Figure 3). Aged mice overexpressing the AMPAR binding domain of NPTX2 (NPTX2-PTX)^18^ in *N*-RAM ensemble, but not in *F*-RAM ensemble rescued overgeneralization in context C during retrieval (Figure 7P-7V, Figure S14E-14F). However, overexpression of NPTX1 and NPTX2-PTX in young mice had no effect on freezing levels when tested in either context A or context C (Figure S15). These data confirm that DG engram network hyperactivity caused by NPTXs downregulation underlies aging-associated memory imprecision, and that re-stabilization of NPTX1-dependent MEC-*Fos^+^*engram excitatory circuit and NPTX2-dependent DG PV^+^ interneuron-*Npas4^+^* engram inhibitory circuit is able to repair memory deficits in aged mice.

**Figure 7.**
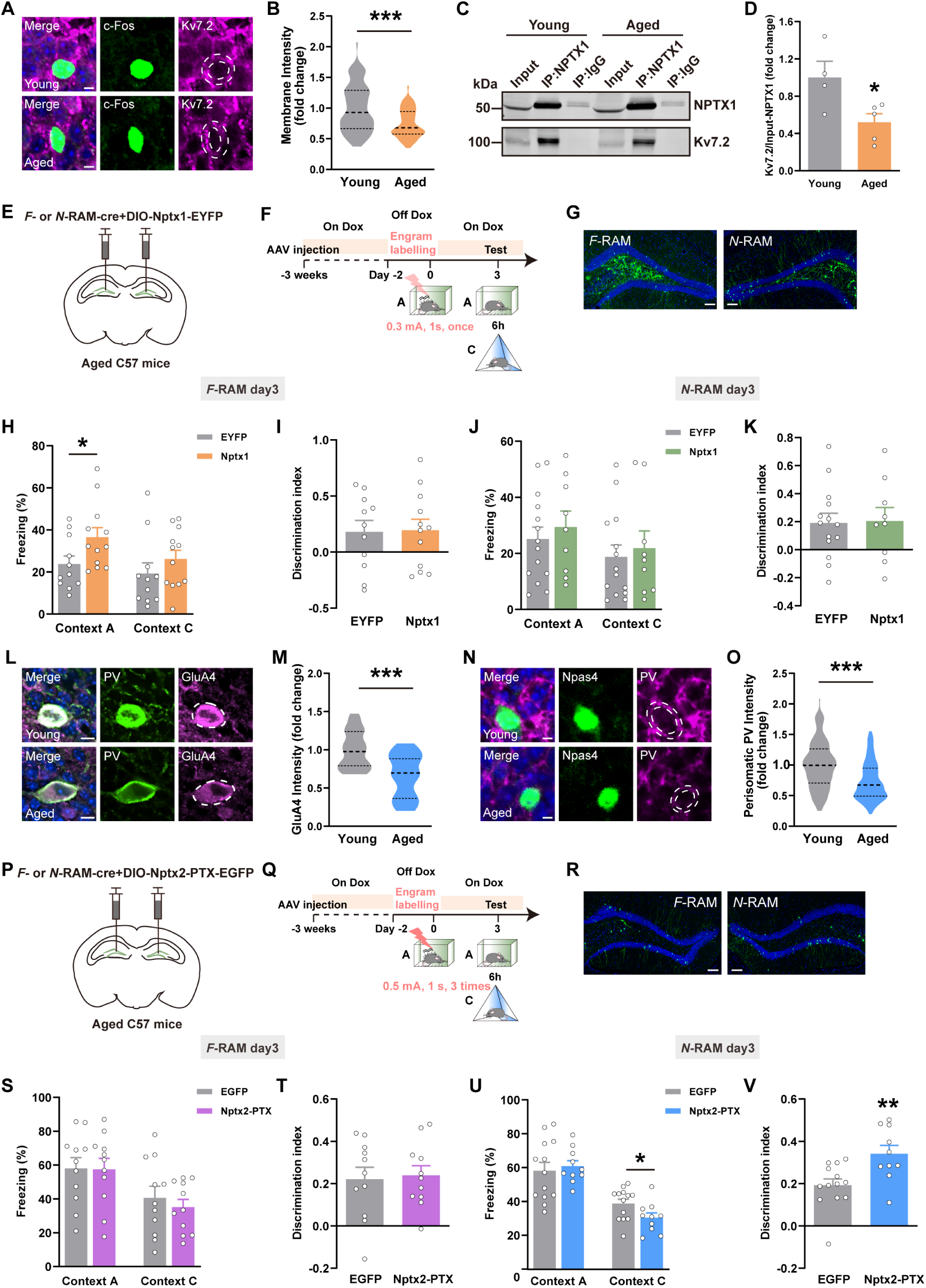
Overexpression of NPTXs in DG engram ensembles rescued memory imprecision in aged mice. (A) Representative confocal images of c-Fos^+^ cells colocalizing with Kv7.2. Green: c-Fos^+^ engram cells, Purple: Kv7.2, Blue: DAPI. Dashed white lines outline cell membrane. Scale bar: 5 μm. (B) The average membrane Kv7.2 fluorescence intensity of c-Fos^+^ neurons in young and aged mice. (C) Immunoblotting of NPTX1 co-immunoprecipitates with Kv7.2 in DG of young and aged mice. (D) The quantification of IP-Kv7.2 in young and aged mice. (E, P) Diagram of AAV injection. (F, Q) Experimental scheme of memory retrieval test. (G, R) Representative expression of NPTX1 or NPTX2-PTX in *F*-or *N*-RAM engram cells in DG. Green: *F*-RAM or *N*-RAM ensemble, EYFP or EGFP, Blue: DAPI. Scale bar: 100 μm. (H, I) The freezing percentage and discrimination index of EYFP and Nptx1 aged groups (*F*-RAM). (J, K) The freezing percentage and discrimination index of EYFP and Nptx1 aged groups (*N*-RAM). (L) Representative confocal images of GluA4 colocalizing with PV^+^ interneurons. Green: PV^+^ interneurons, Purple: GluA4. Scale bar: 10 μm. (M) The quantification of GluA4 membrane expression on PV^+^ interneurons in young and aged mice. (N) Representative confocal images of PV neurites around Npas4^+^ engram cells. Green: Npas4^+^ engram cells, Purple: PV, Blue: DAPI. Dashed white lines outline PV neurites. Scale bar: 5 μm. (O) The average fluorescence intensity of PV neurites around Npas4^+^ neurons in young and aged mice. (S, T) The freezing percentage and discrimination index of EGFP and Nptx2-PTX aged groups (*F*-RAM). (U, V) The freezing percentage and discrimination index of EGFP and Nptx2-PTX aged groups (*N*-RAM). Data are presented as mean ± S.E.M; \**P* < 0.05, \*\**P* < 0.01, \*\*\**P* < 0.001.

## Discussion

Contextual learning activates distinct *Fos^+^* and *Npas4^+^* engram ensembles, recruiting enhanced excitatory and inhibitory circuits respectively to modulate memory-guided adaptive behaviors. However, fundamental questions still remain: how the recruited engram network maintains its stability and whether aging-related memory imprecision associates with engram network destabilization. Our study identified NPTXs as the molecular substrates that prevent network hyperactivity and maintain memory precision. Although the NPTXs expression are not specific to *Fos^+^* and *Npas4^+^* engram cells, the functional specificity of NPTXs should be placed into specific engram circuits to be manifested, and the maintenance of circuit stability depends on the stable expression of NPTX1 and NPTX2 specifically. Molecules and circuits, they are interdependent, both are essential for ensuring precise behavioral outputs.

NPTX1 plays a critical role in stabilizing the excitatory synaptic connection between MEC and DG *Fos^+^* engram cells via facilitating Kv7.2 membrane expression-dependent inhibition of neuronal hyperexcitability to promote memory retrieval. Our data demonstrate that NPTX1 interacts with a specific potassium channel Kv7.2 *in vivo*, although it might not be a direct interaction and probably requires other auxiliary proteins such as syntaxin^28^, we provide a possible explanation for NPTX1 to regulate neuronal excitability beyond its classical role in glutamate signaling. Our behavioral output caused by *Nptx1* depletion in *F*-RAM ensemble was inconsistent with the results of Sun *et al*^17^ that chemogenetic activation of *F*-RAM ensemble or optogenetic inhibition of MEC-DG circuit promoted or inhibited memory generalization. That’s probably because chemogenetic or optogenetic manipulation was transient, while *Nptx1* depletion somehow mimics the persistent activation, leading to engram hyperexcitability and decreased the signal-to-noise ratio (SNR) of MEC-*Fos^+^* ensemble excitatory transmission during memory retrieval. Besides, a number of studies support the view that MEC inputs onto DG regulate memory retrieval^29,30^.

NPTX2 stabilizes the inhibitory synaptic connection between DG local PV^+^ interneurons and *N*-RAM ensemble to suppress memory overgeneralization, consistent with Sun *et al*’s report that *N*-RAM ensemble contributed to memory discrimination^17^. Their finding confirmed the involvement of CCK^+^ interneurons, while our finding illustrated the importance of NPTX2 in mediating PV^+^ interneurons plasticity. We speculate that *N*-RAM ensemble-dependent mediation of memory discrimination may require the coordination of both CCK^+^ and PV^+^ GABAergic interneurons as they have been found to exert complementary roles in recruiting perisomatic inhibition^31,32^ while NPTX2 specifically regulates the functional connection between PV^+^ interneurons and *Npas4*^+^ engram cells. NPTX2 was found as a downstream protein of NPAS4^33,34^, which explained the coincidence of the weakened perisomatic inhibition caused by *Nptx2* knockout with the finding of *Npas4* depletion by Elizabeth A *et al*’s ^35^. As a neuronal IEG, Npas4 upregulates perisomatic inhibitory synapses when activated,^34,36^ to scale down the level of network activity in response to neuronal excitation, but the specific mechanism remains to be elucidated. Our study provide the downstream molecular mechanism for Npas4 activation-dependent mediation of memory precision. The specificity of NPTX2 functions in *Npas4^+^* ensemble but not in *Fos^+^*ensemble to regulate memory discrimination can also be supported by the study by Ee-Lynn *et al* that knockdown of *Nptx2* in *Fos^+^* ensemble had no effect on the perisomatic inhibition received from PV^+^ interneurons^37^.

Memory precision changes dynamically across the lifespan, young individuals only form gist-like memories and slowly develop into precise episodic memories as the brain matures, then the ability to retrieve specific information declines with age. Understanding the maladaptive changes in the engram network stability during aging may provide insights to combat aging-induced cognitive deficits. NPTXs are classic synaptic proteins that may also be prognostic biomarkers in cognitive and mental disorders, such as AD and schizophrenia^14,15^. *Nptx2* was also seen as a neuronal IEG. Our smFISH results showed that the total number of *Nptx2^+^* cells and the expression of *Nptx2* in engram cells were both downregulated in DG during aging, whereas the overall downregulation of *Nptx1* was not observed in aged mice, indicating that NPTX2 was more sensitive to aging. That is to say, along with aging, *Nptx2* downregulation-induced disruption of local perisomatic inhibition of DG engram occurs before *Nptx1* downregulation-induced disruption of MEC-DG engram excitatory long-range projection, which probably explains why generalization always occurs before amnesia in aged individuals. The findings of Ramsaran *et al* showed that extracellular perineuronal nets-dependent functional maturation of PV^+^ interneurons promotes sparse engram formation and memory precision during brain development^38^. Our study elucidated that engram network hyperactivity, caused by NPTXs downregulation contributes to aging-related memory imprecision and provided potential synaptic and molecular strategies to mitigate different phenotypes of aging-related memory imprecision, such as amnesia and overgeneralization respectively.

## Data availability

The *F*– and *N*-RAM engram cells RNA-seq data have been submitted to NCBI Gene Expression Omnibus (GEO) under accession number PRJNA1067898.

## Code availability

Custom codes will be provided upon request.

## Acknowledgement

This work was supported by grants from the STI2030-Major Projects (2021ZD0203500 to FFW and LM, 2021ZD0202100 to XL), the Natural Science Foundation of China (32222033 to FFW, 31930046 and 82021002 to LM, 32171041 to XL, 31970543 and 32270660 to QML), the CAMS Innovation Fund for Medical Sciences (2021-I2M-5-009 to LM and XL). We thank Prof. Yingxi Lin (The University of Texas Southwestern Medical Center, Texas, USA) for providing the *AAV-Fos-RAM-d2tTA-TRE-mKate2* and *AAV-Npas4-RAM-d2tTA-TRE-mKate2* plasmids.

## Author contributions

F.W., L.M. and T.J. designed the research. T.J. and Y.Y. conducted the AAV vector construction, virus injection, immunostaining, Co-IP, smFISH, Ribo Tag purification and the behavioral experiments. T.J. carried out the electrophysiology recordings. N.H., Y.G., Y.Z and Q.L. contributed to the bioinformatics analysis. T.J., Y.Y. J.Y, and X.L. analyzed the data and T.J. carried out the statistical analysis and drafted the manuscript. L.M. and F.W. supervised the project and revised the paper.

## Methods

### Animals

All animals used in this study were listed as follows: adult C57BL/6J male mice (3 m) from the SLAC Laboratory Animal Company (Shanghai, China), aged C57BL/6J male mice (16-18 m) from the Aniphe Biolaboratory Inc. (Nanjing, China) and *PV-Flpe* (021191) mice from the Jackson Laboratory (CA, USA). CRISPR-Cas9 mediated construction of *Nptx1^fl/fl^*, *Nptx2 ^fl/fl^* conditional knockout (*Nptx1* cKO, *Nptx2* cKO) mice targeting exon 3 and exon 2 respectively were generated by the Biocytogen Co. (Beijing, China) and *Npas4-CreER^T^*^2^ mice were generated by Shanghai Model Organisms center, Inc (Shanghai, China). *Npas4-CreER^T^*^2^ and *PV-Flpe* mice were bred to C57BL/6J mice for more than six generations, *Nptx1^fl/fl^*, *Nptx2 ^fl/fl^* mice and their respective wild-type (WT) littermates were obtained from self-crossing of *Nptx1^fl/+^*, *Nptx2 ^fl/+^* (heterozygous) mice. Genotypes were determined by polymerase chain reaction (PCR) of mouse toe DNA samples. The primers for genotyping PCR were as follows: 5’ – GCTGTAGGGATGCTTGTCTCTGGTG – 3’ and 5’ – AGAAAAGCTGACCCAAGGTCTCTGC – 3’ for Nptx1 5’ loxP, 5’ – ATTAGCTGCCAGATCTTAGCCCCCT – 3’ and 5’ – GTGTGTGTCCCTGGTGGTGAAGTTT – 3’ for Nptx1 3’ loxP, 5’ – GAATGGCTCGAGGCAGGTCCAGTTT – 3’ and 5’ – CGTTACTAAACCCCAGACAGCTCCG – 3’ for Nptx2 5’ loxP, 5’ – AGTTCTGCCTCTGTTCATCTTGCCA – 3’ and 5’ – TTCACCTGACCCTTCTGTTCACGAC – 3’ for Nptx2 3’ loxP, 5’ – AGAGCCTGAGCGAAAAGACC – 3’, 5’ – CTGCTCACCTCCAGCAAAGA – 3’ and 5’ – CGCGCGCCTGAAGATATAGA – 3’ for *Npas4-CreER^T^*^2^. Male offsprings at 8-12 weeks of age were used in following experiments, which were randomly assigned to groups. All mice were housed on a 12 hr light/dark cycle (light on from 8 a.m. to 8 p.m.) with access to food and water *ad libitum*. All experiment procedures were strictly in accordance with the National Institutes of Health Guide for the Care and Use of Laboratory Animals, and were approved by Animal Care and Use Committee of the animal facility at Fudan University.

### Viral vectors

*AAV-Fos-RAM-d2tTA-TRE-mKate2* and *AAV-Npas4-RAM-d2tTA-TRE-mKate2* plasmids were kind gifts from Prof. Yingxi Lin (The University of Texas Southwestern Medical Center, Texas, USA). To generate *AAV-Fos-RAM-d2tTA-TRE-Cre* and *AAV-Npas4-RAM-d2tTA-TRE-Cre* plasmids, *mKate2* in *AAV-Fos-RAM-d2tTA-TRE-mKate2* and *AAV-Npas4-RAM-d2tTA-TRE-mKate2* plasmids was replaced with the *Cre* sequence obtained by PCR from *pAAV-Cre-GFP* (Addgene: 68544). *pAAV-CMV-bGlobin-Flex-EGFP-MIR30-Scramble-shRNA*, *pAAV-CMV-bGlobin-Flex-EGFP-MIR30-Nptx2-shRNA* and *pAAV-CMV-DIO-Nptx2-PTX-P2A-EGFP* were used in our previous paper^18^. To generate *pAAV-Ef1α-DIO-Nptx1-T2A-EYFP*, the coding sequence for mouse *Nptx1* with *T2A* was generated by Azenta US, Inc. (Suzhou, China) and was subcloned into *pAAV-Ef1α-DIO-EYFP* (Addgene: 27056). Adeno-associated viruses (AAVs) described above were packaged by OBiO Technology Co., Ltd (Shanghai, China) into serotype 9. *AAV*_9_*-Ef1α-Flex-NBL10* (69971) and *AAV*_9_*-Ef1α-DIO-ChR2-EYFP* (S0199-9-H50) were purchased from Taitool Bioscience Co., Ltd. (Shanghai, China). *AAV*_9_*-Ef1α-DIO-H2B-EGFP* (PT-0258), *AAV*_9_*-Ef1α-DIO-EYFP* (PT-0012), *AAV*_9_*-CaMKIIα-ChR2-mCherry* (PT-0297), *AAV*_9_*-VGAT2-Flp* (PT-2501, vector backbone from BrainVTA)*, AAV*_9_*-CCK-fDIO-mCherry* (PT-8508, vector backbone from Addgene: 114471)*, AAV*_8_*-Ef1α-DIO-TVA-H2B-EGFP* (PT-0021), *AAV*_8_*-Ef1α-DIO-RVG* (PT-0023) and *RV-ENVA-ΔG-dsRed* (R01002) were purchased from BrainVTA Co., Ltd. (Wuhan, China). *AAV*_9_*-PV-Flp* (BC-0430, vector backbone from Addgene: 22914), *AAV*_9_*-SST-Flp* (BC-0429, vector backbone from Addgene: 22913)*, AAV*_9_*-Ef1α-fDIO-ChR2-mCherry* (BC-0113), *AAV*_9_*-Ef1α-fDIO-mCherry* (BC-0193) and *AAV*_9_*-Ef1α-fDIO-hM3Dq-mCherry* (BC-0495) were purchased from Brain Case Co., Ltd. (Shenzhen, China).

### Stereotaxic surgery

Mice were anesthetized with 2% isoflurane and placed in a stereotaxic instrument (RWD Life Science, Shenzhen, China). Microinjections were performed using 33-gauge needles connected to a 10 μl microsyringe (Hamilton, Bonaduz, Switzerland), which were under the control of a UMP3 ultra micropump (World Precision Instruments, Florida, USA). The coordinates relative to bregma were listed as follows: anterior-posterior (AP) – 1.9 mm; medial-lateral (ML) ± 1.1 mm; dorsal-ventral (DV) – 2.1 mm for DG and AP – 4.7 mm; ML ± 3.2 mm; DV – 4.5 mm for MEC. The needle was slowly lowered to the target site and remained for at least 3 min after injection. The injection volume per site was 0.3 μl in DG and 0.4 μl in MEC. The final titer of all the AAVs used for infections were at least 1 × 10^12^ V.G./ml except for *AAV-SST-Flp, AAV-PV-Flp, AAV-VGAT2-Flp, AAV-Fos-RAM-d2tTA-TRE-Cre* and *AAV-Npas4-RAM-d2tTA-TRE-Cre,* which were 1:1000 diluted. For rabies infections, the final titer used was 1 × 10^8^ IFU/ml. After surgery, all mice were given at least 3 weeks to recover before behavioral experiments or electrophysiological recordings, and the efficiency of virus infection was verified by immunostaining. Only the mice with virus infection in correct places were chosen for further analysis.

### Engram labeling

To label engram cells, *Fos-RAM-d2tTA-TRE (F-RAM)* and *Npas4-RAM-d2tTA-TRE (N-RAM)* systems, which gene expression was under the control of the tetracycline responsive element (TRE), were used respectively through AAVs infusion. All mice were kept on doxycycline (Dox, Huamaike Bio, 360304) containing food (40 mg/kg) one day before virus injection. 48 hr (day –2) before engram labeling, mice were fed with regular food instead of Dox diet. Then contextual fear conditioning or novel context exposure (context N) was performed on day 0 to label *Fos^+^* or *Npas4^+^* engram cells in DG, after labeling, mice were put on Dox containing food again immediately. Mice were given at least 3 days to allow sufficient protein expression before subsequent experiments.

### Rabies input tracing

To study the respective monosynaptic inputs of *Fos^+^*and *Npas4^+^* ensembles in DG, Rabies trans-synaptic tracing experiment was performed. Mice were infected with *AAV*_9_*-Fos-RAM-d2tTA-TRE-Cre* or *AAV*_9_*-Npas4-RAM-d2tTA-TRE-Cre,* AAV helper (*AAV*_8_*-Ef1α-DIO-TVA-H2B-EGFP* and *AAV*_8_*-Ef1α-DIO-RVG*). On day 3 after engram labeling, Rabies virus (*RV-ENVA-ΔG-dsRed*) was injected into DG at the same coordinates, then mice were housed in the BSL2 facility for 1 week to allow rabies spread and dsRed expression before perfusion (day 10). For rabies tracing analysis, consecutive brain slices from AP + 2.8 mm to AP – 5.0 mm (40 μm thickness) selected from every fifth slice were collected. DsRed^+^ cells were manually counted by an experimenter who was blind to the experimental condition. The percentage of rabies-labelled inputs was calculated as follows, dsRed^+^ cells in each brain region / total dsRed^+^ cells per mouse.

### Contextual fear conditioning (CFC)

Before CFC was performed, mice were handled daily in a holding room for 3 days. For experiments with Dox-dependent ensembles labeling, on the third handling day, the Dox diet was replaced with regular food (off-Dox) and CFC assay was typically carried out 48 hours after the last handling session. CFC was performed in the conditioning chamber (Med-Associates, St. Albans, VT, USA) and the procedure was composed of conditioning and test sessions.

On the conditioning day (day 0), mice were firstly transported into the holding room and allowed to habituate for at least 30 minutes, then transported into the behavioral room and placed into context A, a square plexiglass observational chamber with stainless steel bars connected to a shock generator on the floor for conditioning. Two individual conditioning protocols were used in our study, for mice that conditioned to the 360 s protocol, three foot shocks (0.5 mA, 1 s each) at 180 s, 240 s and 300 s were given and mice were taken out 60 s after termination of the third foot shock, for mice that conditioned to the180 s protocol, one single foot shock (0.3 mA, 1 s) at 120 s was given and mice were taken out 60 s after termination of the foot shock.

The test session was carried out on day 3. Mice were put back into fear context A for 180 s, and were placed into a non-fear context (context C, a triangular chamber with white, smooth plastic floor and black cover) for 180 s 6 hr later. For immunohistochemistry, mice were tested for memory expression in only one context, either context A or C, and sacrificed 1 hr later.

The freezing percentage was automatically analyzed by software (Med-Associates) with freezing defined as absence of movement for at least 1 s. The discrimination index (DI) was calculated as follows, (the freezing percentage in context A – the freezing percentage in context C) / (the freezing percentage in context A + the freezing percentage in context C).

### Open field test (OFT)

Spontaneous locomotor activity was carried out as we previously reported^39^, in short, mice were placed in the center of an open arena (40 × 40 cm^2^) at the beginning of the test, and were allowed to freely explore the arena for 20 min. Distance travelled in the arena and time in center zone were quantified using a TopScan automated detection system (CleverSys, Reston, VA, USA).

### O-maze test

The O-maze was 70 cm in diameter and 70 cm high off the ground, and consisted of two open arms (7 cm wide) without walls, two closed arms that were enclosed by vertical walls. Mice were gently placed into the open arms, and their behavior was recorded for 6 min via a TopScan automated detection system (CleverSys, Reston, VA, USA) located above the maze.

### Tail suspension test (TST)

Mice were suspended 20 cm above a solid surface by the use of adhesive tape applied to the tail, and their behavior was recorded for 6 min. Immobility time was defined as absence of struggling for at least 1 s and latency to immobility was defined as the time from the beginning of TST to mice’s first absence of struggling for at least 1 s, which were manually analyzed by an experimenter who was blind to the experimental condition.

### Drug injections

Clozapine N-oxide (CNO, Sigma, C0832) was dissolved in saline, which was intraperitoneal injected (i.p.) at a concentration of 1 mg/kg 30 min before memory retrieval. The KCNQ2/3 activator, retigabine (Supelco, 90221) was diluted in 5% DMSO (Sigma, 276855) and administered i.p. at 1 mg/kg 30 min before memory retrieval with 5% DMSO as control. 4-Hydroxytamoxifen (4-OHT, Sigma, H6278) was dissolved in ethanol at 20 mg/ml and stored at –20℃. The ethanol was evaporated at 95℃ and corn oil (Thermo Fisher, C40543) was added to give a final concentration of 10 mg/ml, which was i.p. injected 2 hr before memory conditioning with corn oil as control.

### Whole-cell patch-clamp recording in brain slices

Mice were subjected to electrophysiology on day 3 following CFC. Living acute brain slices (300 μm) preparation for analyzing engram synaptic connections and engram intrinsic excitability were performed as previously described^40^ only with minor modifications. Briefly, mice’s brains were cut on a vibratome (Thermo Scientific, MA, USA) in carbogenated (95% O_2_, 5% CO_2_) ice-cold cutting solution containing (in mM): 93 NMDG, 2.5 KCl, 1.25 NaH_2_PO_4_, 30 NaHCO_3_, 20 HEPES, 25 glucose, 2 thiourea, 5 Na-ascorbate, 3 Na-pyruvate, 0.5 CaCl_2_ and 10 MgCl_2_, 300-310 mOsm, pH adjusted to 7.3 with HCl. After initial recovery at 32 °C for 10 min, slices were transferred to carbogenated HEPES holding ACSF (in mM: 92 NaCl, 2.5 KCl, 1.25 NaH_2_PO_4_, 30 NaHCO_3_, 20 HEPES, 25 glucose, 2 thiourea, 5 Na-ascorbate, 3 Na-pyruvate, 2 CaCl_2_ and 2 MgCl_2_, 300-310 mOsm, pH 7.4) and incubated for over 45 min before recording. Whole-cell patch-clamp recordings were performed in carbogenated recording ACSF (in mM: 119 NaCl, 2.5 KCl, 1.25 NaH_2_PO_4_, 24 NaHCO_3_, 12.5 glucose, 2 CaCl_2_ and 2 MgCl_2_ (300-310 mOsm, pH 7.3-7.4) at a rate of 1.5 mL / min (30∼32°C) with an EPC-10 amplifier and Pulse v8.78 software (HEKA Elektronik, Lambrecht/Pfalz, Germany). Recording neurons were identified visually by location, morphology, size and fluorescence and recordings were performed using borosilicate glass pipettes (5-7 MΩ tip resistance).

To record A/N ratios, neurons were voltage-clamped at –70 mV to record AMPAR mediated EPSCs and at +40 mV to record dual-component EPSCs containing NMDA receptor (NMDAR) EPSCs. To calculate the A/N ratios, the peak current of the AMPA EPSC at –70 mV was compared with the value of the NMDA EPSC after stimulation start time 50 ms at +40 mV. Intracellular solution used was (in mM): 127.5 cesium methanesulfonate, 7.5 CsCl, 10 HEPES, 2.5 MgCl_2_, 4 Mg-ATP, 0.4 Na_3_-GTP, 10 sodium phosphocreatine, 0.6 EGTA (290-300 mOsm, pH 7.2).

To record light-evoked EPSCs (oEPSC), A TTL-driven light-emitting diode (Lumen Dynamics) was used to generate photostimulation consisting of a single wide-field blue flash (470 nm, 1 ms duration) for photostimulation of ChR2-expressing terminals. The laser intensity was measured at the focal plane of the slice when delivered through the 40× water-immersion objective lens (Nikon, Japan). Slices containing MEC ChR2-expressing terminals were chosen for oEPSC recording in the presence of 1 μM Tetrodotoxin (TTX, Tocris, 1078), 100 μM 4-aminopyridine (4-AP, Sigma, 275875) and 100 μM picrotoxin (PTX, Tocris, 1128), neurons were voltage-clamped at –70 mV and the stimulus intensity was 0.1 mW at 470 nm. To record paired-pulse ratio (PPR) and AMPA/NMDA (A/N) EPSC ratio, DG slices containing ChR2-expressing neurons were chosen to perform patch-clamp recordings in the presence of 100 μM PTX and the stimulus intensity was 0.1 mW at 470 nm. The AMPA receptor (AMPAR) mediated EPSCs were evoked by paired photostimulation of 50 ms interval for 10 consecutive traces, and PPR was determined as the peak amplitude ratio of the second to the first EPSC. To record light-evoked IPSCs (oIPSC) and PPR, neurons were clamped at +10 mV in the presence of 1 μM TTX, 100 μM 4-AP, 20 μM CNQX (Sigma, C127) and 50 μM D-AP5 (Tocris, 0106) and the stimulus duration and intensity was 1 ms, 0.2 mW at 470 nm. The recording protocols and intracellular solution used were the same as above. To record Action Potentials (AP), current-clamp was used and membrane potentials were measured in response to intracellular injection of step currents (1000 ms duration, magnitudes ranging from – 150 to 250 pA in steps of 10 pA) with the addition of 20 μM CNQX, 50 μM D-AP5 and 100 μM PTX into ACSF. Intracellular solution used was (in mM): 135 potassium-gluconate, 4 KCl, 2 NaCl, 10 HEPES, 4 EGTA, 4 Mg-ATP, 0.3 Na3-GTP, 10 sodium phosphocreatine (280-290 mOsm, pH 7.3).

To record and isolate M-current (*I*_M_), intracellular solution used was the same as above, 0.2 mM CdCl_2_ (Sigma, 202908), 1 μM TTX, 10 μM ZD7288 (Tocris, 1000), and 4 mM 4-AP to block voltage-dependent Cav, Nav, HCN and Kv1 channels, 20 μM CNQX, 50 μM D-AP5 and 100 μM picrotoxin to block synaptic activity were added into ACSF. To isolate M-current, the following protocol the same as previously reported^41^ with minor modifications was applied to the recording cells: (1) a 1 s step from the holding potential (–70 mV) to –10 mV was applied to activate *I*_M_ while inactivating most other voltage-gated currents, (2) a 1 s step from –60 mV to –10 mV with 10 mV increments to elicit *I*_M_ tail current, and (3) a 1 s step to –10 mV, before returning to –70 mV.

The signals were acquired at 10 kHz and filtered at 2 kHz. The series resistance was < 30 MΩ. Data were analyzed with Mini Analysis Program (Synaptosoft, Fort Lee, NJ, USA) or pCLAMP10.7 (Molecular Devices, San Jose, CA, USA) by an experimenter who was blind to the experimental condition.

### Ribosome-Associated Messenger RNA Purification/ RiboTag

DG from mice injected *AAV*_9_*-F/N-RAM-Cre* and *AAV*_9_*-Flex-NBL10,* Cre-dependent expression of N terminus of ribosomal subunit protein Rpl10a (NBL10)^42^ following fear conditioning (Day 3) was quickly isolated and used for enrichment of ribosome-associated transcripts as described previously^18^. The brain tissue (DG) was homogenized in supplemented hybridization buffer (HB-S) containing dithiothreitol (DTT, Sigma, D9760), cycloheximide (CHX, Cayman, 14126), heparin (Tocris, 2812), protease inhibitors (Roche, 04693116001), and RNase inhibitor (Vazyme, R301). The supernatant was incubated with anti-hemagglutinin (HA) antibody (Sigma, H6908) and Dynabeads Protein G (Invitrogen, 10003D) for 12 hours. Purified messenger RNA (mRNA) was eluted from the Dynabeads using TRIzol^TM^ LS (Invitrogen, 10296010). An Agilent RNA 6000 Pico Kit (Agilent, 5067-1513) and an Agilent 2100 bioanalyzer were used to evaluate the quality of purified mRNA. Purified mRNA samples with RNA integrity number < 7 were discarded.

### Sequence Processing and Data Analysis

The library was prepared with VAHTSTM mRNA-seq V3 Library Prep Kit for Illumina (Vazyme, NR611) and sequenced on a HiSeq 4000 (Illumina) by Novogene Technology Co., Ltd (Beijing, China). Raw reads were cleaned with FASTX-toolkit to remove adapter contamination and low-quality reads (quality score < 28). The clipped reads were aligned to mouse reference sequence (GRCm38/mm10) using HISAT2^43^. Cuffdiff-generated FPKM count matrix was used for subsequent analysis. Significance was drawn with ANOVA and clustered using Shannon entropy-based method^44^. Genes with more than one and half fold expression changes, and were significantly different (*P* < 0.05) were selected for further analysis.

### Single molecule fluorescent in situ hybridization (smFISH)

Fixed-frozen brain tissues were sliced into 10 μm coronal sections and baked at 60°C for 30 min. Then slices were incubated with hydrogen peroxide (H_2_O_2_) for 10 min at room temperature (RT), targets retrieval and protease III incubation were performed using RNAscope® 2.5 Universal Pretreatment Reagents (Advanced Cell Diagnostics, 322000, 322381). SmFISH probes for all genes examined: *Nptx1* (505421), *Nptx2* (316901), *Fos* (316921), *Npas4* (423431), *Sirt1* (418341) and *Atg7* (561261) were hybridized for 2 hr. After Hybridization, RNAscope® Multiplex Fluorescent Detection Kit v2 (Advanced Cell Diagnostics, 323110) were used to amplify signals. Images were acquired by using the Nikon A1 confocal microscope (Tokyo, Japan). Regions of interest (ROI) were circled and the intensity within the regions was analyzed by Image-Pro Plus 6.0 (Media Cybernetics, Rockville, MD, USA). For cell counting analysis, consecutive brain slices from AP – 1.2 mm to AP – 2.0 mm (10 μm thickness) selected from every tenth slice, 8 slices per mouse were collected, number of mice were included in the statistical table.

### Immunohistochemistry (IHC)

Mice were perfused transcardially with ice cold saline followed by 4% paraformaldehyde (PFA, dissolved in 0.1 M PBS). The brains were removed and fixed in 4% PFA overnight. Then the brains were subjected to dehydrate in 30% sucrose solutions at 4°C for 72 hr before being sliced into 30 or 40 μm coronal sections (selected from every fifth slice). Slices were incubated with primary antibodies in blocking solution containing 0.3% Triton X-100 overnight at 4°C. Slices were washed with 0.1 M PBS, and then incubated with secondary antibody at RT for 1.5 hr. After being washed in PBS, slices were mounted in anti-quenching mounting medium (Southern Biotech, 0100). Primary antibodies used were: anti-NPTX1 (1:400, Alomone Labs, ANR-191), anti-NPTX2 (1:500, Proteintech, 10889-1-AP), anti-Fos (1:500, Synaptic Systems, 226017), anti-Parvalbumin (PV, 1:100, Thermo Fisher, PA1-933), anti-PV (1:500, Oasis, OB-PGP005-01), anti-GluA4 (1:500, Merck, AB1508), anti-Somatostatin (SST, 1:200, Oasis, OB-PRB111-01), anti-Choline Acetyltransferase (ChAT, 1:100, Merck, AB143), anti-Glutamate (1:500, Merck, AB133), anti-GAD67 (1:500, Merck, MAB5406), anti-cholecystokinin (CCK, 1:1000, Merck, C2581) and anti-KCNQ2 (1:100, Alomone Labs, APC-050). Secondary antibodies used were: goat anti-rat 488 (1:1000, Jackson Immuno Research, 112-545-167), goat anti-rabbit 488 (1:1000, Jackson Immuno Research, 111-545-144), donkey anti-guinea pig 488 (1:1000, Jackson Immuno Research, 706-545-148), goat anti-rabbit Cy3 (1:1000, Jackson Immuno Research, 111-165-144), donkey anti-rat Cy3 (1:1000, Jackson Immuno Research, 712-165-153) and donkey anti-rat 647 (1:500, Jackson Immuno Research, 712-605-150). Kv7.2 and GluA4 membrane expression analysis were adapted from one of the latest researches from our lab^45^. Images were acquired using a Nikon-A1 confocal microscope (Tokyo, Japan) with a 20× objective lens or a 60× objective oil lens. Data were analyzed blindly to the group using Image-Pro Plus 6.0 and ImageJ (Fiji). For cell counting analysis, consecutive brain slices from AP – 1.2 mm to AP – 2.0 mm (40 μm thickness) selected from every fifth slice, 4 slices per mouse were collected, number of mice were included in the statistical table.

### Reverse transcription-quantitative PCR (RT-qPCR)

Reverse transcription was completed using HiScript II 1st Strand cDNA Synthesis Kit (Vazyme, R212). The cDNA was subjected to qPCR using ChamQ Universal SYBR qPCR Master Mix (Vazyme, Q711) and Eppendorf Mastercycler PCR System. The primers for qPCR were as follows: 5’ – ACCTCCCTACACCAACGGAT – 3’ and 5’ – GGCAGGCTCTTCTTCACCTT – 3’ for *Nptx1*, 5’ – GCCAAGGTGAAGAAGAGCCT – 3’ and 5’ – AGCATAAGAGAAGGGTGTGCC – 3’ for *Nptx1* of exon 3, 5’ – GACTTCCGAGAGGTGCTCCA – 3’ and 5’ – GGTGAGCCGAGGTCTCATTA – 3’ for *Nptx2* of exon 2, 5’ – TGGCCTTCCGTGTTCCTAC – 3’ and 5’ – GAGTTGCTGTTGAAGTCGCA – 3’ for *Gapdh*, which were synthesized by Azenta US, Inc.. The mRNA expression of *Nptx1* and *Nptx2* was normalized to the internal control *Gapdh*.

### Co-immunoprecipitation (Co-IP) and Western blotting

The DG tissues of adult C57BL/6J male mice were rapidly extracted and homogenized in radio immunoprecipitation assay (RIPA) buffer containing protease and phosphatase inhibitors for 30 min. The protein lysate concentration was determined using bicinchoninic acid kit. Protein samples were incubated with 3 µg anti-NPTX1 (Santa Cruz Biotechnology, sc-374199) or anti-IgG (Cell Signaling Technology, 5415S) for 4 h at 4°C. Pierce Protein A/G magnetic beads (88803, Thermo Scientific) were added to each sample and the mixture was rotated at 4°C overnight. The attached protein lysis was then eluted and used for further protein blotting assay. Before Western blotting assay, 50 µg protein samples were boiled for 6 min at 85°C, and were separated by SDS-polyacrylamide gel electrophoresis (PAGE) under 120 V for 1.5 h. Proteins were then transferred to the polyvinylidene fluoride (PVDF) membrane at 100 mV for 100 min. The membrane was firstly washed with Tris-buffered saline with Tween 20 (TBST), followed by blocking in 5% skimmed milk for 2 hours. The membrane was then incubated with the following primary antibodies anti-NPTX1 (1:100, Santa Cruz Biotechnology, sc-374199) and anti-KCNQ2 (1:500, Abcam, ab22897) at 4°C overnight. On the next day, the membrane was washed with TBST and incubated with IRDye 700DX-or 800DX-conjugated anti-rabbit or anti-mouse IgG (1:50000, Rockland Immunochemicals Inc.) for 2 h at room temperature. Protein bands were visualized using Odyssey (LI-COR Biosciences). The immunoblots were analyzed with Image J.

### Statistics

All data were presented as mean ± SEM. Sample sizes were based on our previous research^18,39,46^. Statistical analyses were performed by SPSS 20.0 software (IBM, Armonk, NY, USA) or R software (version 4.1.2). The normality test of the data sets was performed by the Shapiro-Wilk test or the Kolmogorov-Smirnov test if n > 50, homogeneity of variance test of the data sets was performed by the Levene’s test. Two-tailed unpaired *t* test was used for comparing two independent groups. Multiple group comparisons were assessed using One-way analysis of variance (ANOVA), Two-way ANOVA or Two-way repeated measures (RM) ANOVA, followed by the Bonferroni’s post-hoc test when significant main effects or interactions were detected. Mann-Whitney *U* test or Kruskal-Wallis H test was used when normality was violated. Bonferroni’s corrections were applied to assess statistical significance.* *P* < 0.05, ** *P* < 0.01, and *** *P* < 0.001.

