## Supplemental figures and figure legends for "Disturbed engram network caused by NPTXs downregulation underlies aging-related memory deficits"

**This file includes:**

**Supplemental figures: S1-S15**

**Supplemental figure legends**

### Fig. S1

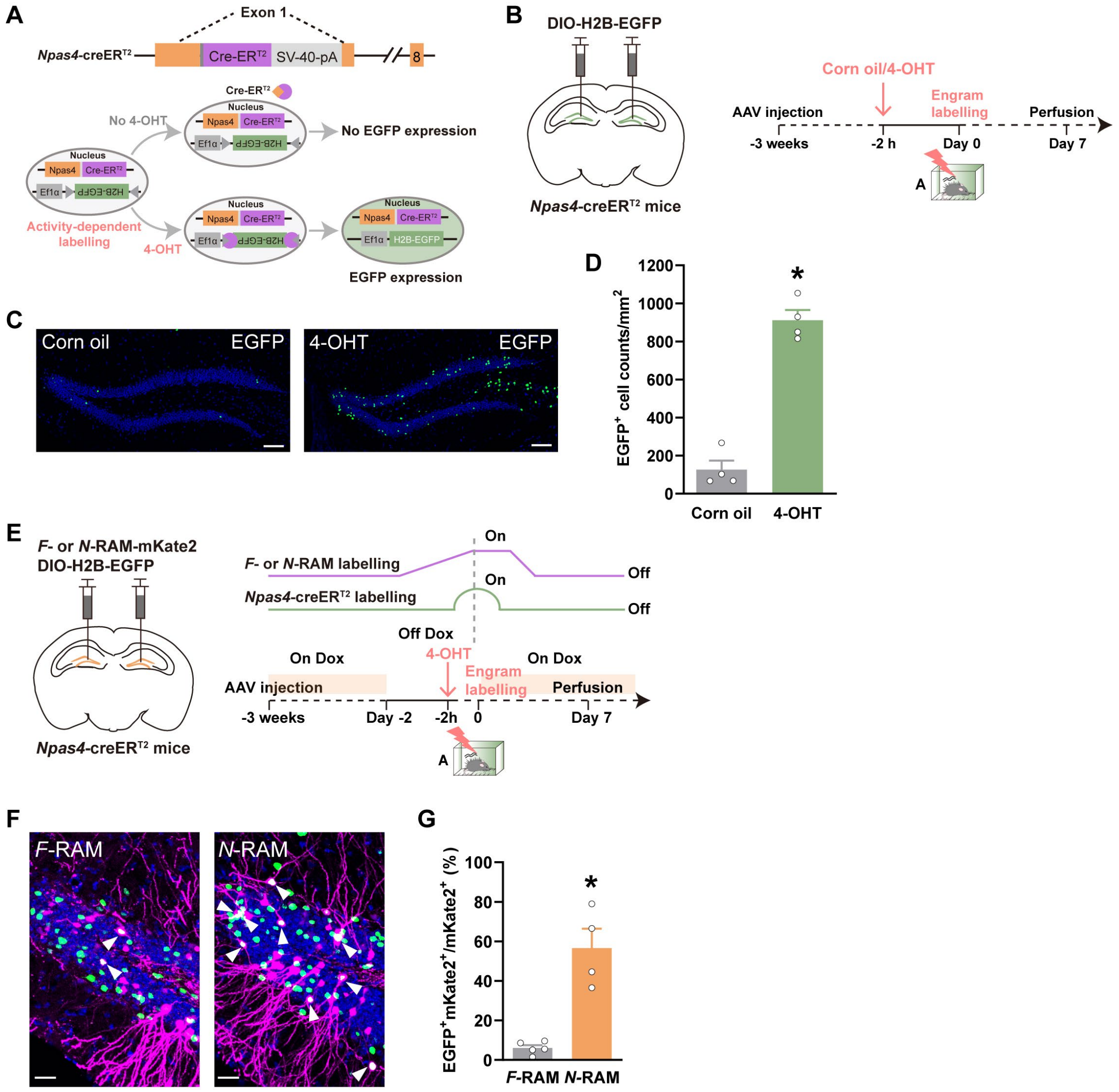
A
Exon 1
Npas4-creER^T2
Cre-ER^T2
SV-40-pA
8
No 4-OHT
Nucleus
Npas4
Ef1α
H2B-EGFP
No EGFP expression
Activity-dependent labelling
4-OHT
EGFP expression
B
DIO-H2B-EGFP
Npas4-creER^T2 mice
Corn oil/4-OHT
AAV injection
Engram labelling
Perfusion
-3 weeks
-2 h
Day 0
Day 7
C
Corn oil
EGFP
4-OHT
EGFP
D
EGFP+ cell counts/mm2
1200
1000
800
600
400
200
0
Corn oil
4-OHT
E
F- or N-RAM-mKate2
DIO-H2B-EGFP
Npas4-creER^T2 mice
F- or N-RAM labelling
Npas4-creER^T2 labelling
On
Off
Off Dox
On Dox
4-OHT
Engram labelling
AAV injection
Perfusion
-3 weeks
Day -2
-2h
0
Day 7
F
F-RAM
N-RAM
G
EGFP+mKate2+/mKate2+ (%)
100
80
60
40
20
0
F-RAM
N-RAM

**Figure S1. Generation of *Npas4-CreER$^{T2}$* mice and the overlapping analysis of *F*-RAM and *N*-RAM engram cells in DG.** (A) Diagram of the targeting strategy for generating *Npas4-CreER$^{T2}$* mice. (B) Diagram of AAV injection and experimental scheme to label *Npas4$^+$* neurons. (C, D) Representative confocal images and quantification of *Npas4$^+$* cells with or without 4-OHT injection. Green: *Npas4$^+$* engram cells, EGFP. Scale bar: 100 μm. (E) Diagram of AAV injection and experimental scheme to label *Npas4$^+$* and *F*- or *N*-RAM neurons. (F, G) Representative confocal images and overlapping analysis of *Npas4$^+$* and *F*-or *N*-RAM cells. Green: *Npas4$^+$* engram cells, EGFP, purple, *F*- or *N*-RAM cells, mKate2, blue, DAPI. White arrows indicate the colocalized neurons. Scale bar: 30 μm. Data are presented as mean ± S.E.M; *$P < 0.05$.

### Fig. S2

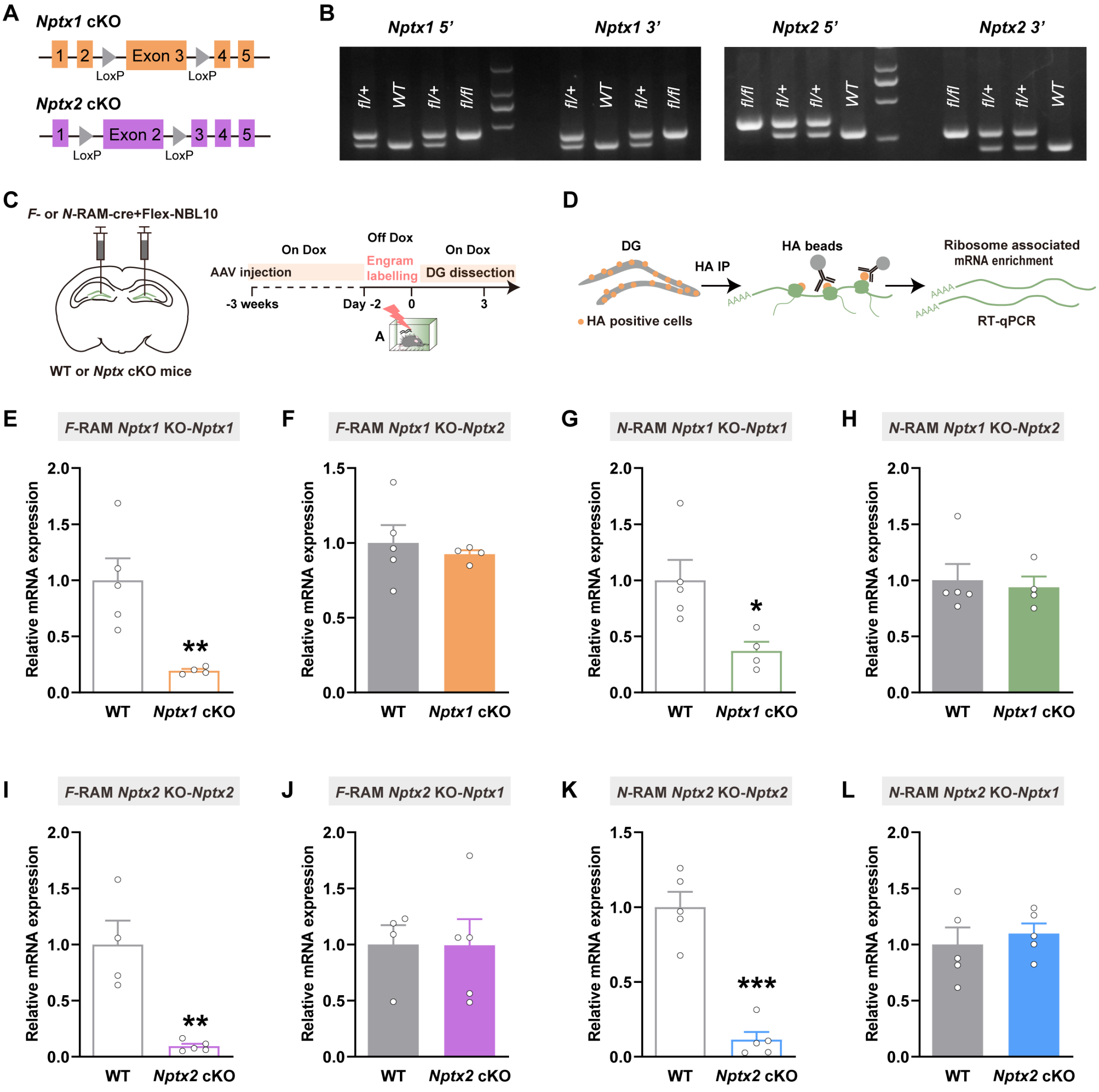
A
Nptx1 cKO
LoxP
Nptx2 cKO
B
Nptx1 5'
Nptx1 3'
Nptx2 5'
Nptx2 3'
C
F- or N-RAM-cre+Flex-NBL10
WT or Nptx cKO mice
AAV injection
On Dox
Off Dox
Engram labelling
On Dox
DG dissection
-3 weeks
Day -2
0
3
D
DG
HA IP
HA beads
Ribosome associated mRNA enrichment
HA positive cells
RT-qPCR
E F-RAM Nptx1 KO-Nptx1
F F-RAM Nptx1 KO-Nptx2
G N-RAM Nptx1 KO-Nptx1
H N-RAM Nptx1 KO-Nptx2
I F-RAM Nptx2 KO-Nptx2
J F-RAM Nptx2 KO-Nptx1
K N-RAM Nptx2 KO-Nptx2
L N-RAM Nptx2 KO-Nptx1
Relative mRNA expression
WT
Nptx1 cKO
Nptx2 cKO

**Figure S2. Generation of *Nptx1* cKO and *Nptx2* cKO mice and RT-qPCR validation in DG engrams.** (A) Diagram of the targeting strategy for generating *Nptx1* cKO and *Nptx2* cKO mice. (B) Genotyping PCR results for $Nptx1^{+/+}$ (*Nptx1* WT), $Nptx1^{fl/+}$ (*Nptx1* heterozygote) $Nptx1^{fl/fl}$ (*Nptx1* cKO), $Nptx2^{+/+}$ (*Nptx2* WT), $Nptx2^{fl/+}$ (*Nptx2* heterozygote) $Nptx2^{fl/fl}$ and (*Nptx2* cKO) mice. (C) Diagram of AAV injection and experimental scheme to label *F*-RAM and *N*-RAM ensembles in WT and *Nptxs* cKO mice. (D) Scheme of RiboTag enrichment of *F*- and *N*-RAM transcriptomes. (E, F) RT-qPCR analysis of *Nptx1* exon 3 and *Nptx2* exon 2 mRNA expression in *F*-RAM ensemble of WT and *Nptx1* cKO mice. (G, H) RT-qPCR analysis of *Nptx1* exon 3 and *Nptx2* exon 2 mRNA expression in *N*-RAM ensemble of WT and *Nptx1* cKO mice. (I, J) RT-qPCR analysis of *Nptx2* exon 2 and *Nptx1* exon 3 mRNA expression in *F*-RAM ensemble of WT and *Nptx2* cKO mice. (K, L) RT-qPCR analysis of *Nptx2* exon 2 and *Nptx1* exon 3 mRNA expression in *N*-RAM ensemble of WT and *Nptx2* cKO mice. Data are presented as mean ± S.E.M; *$P < 0.05$, **$P < 0.01$, ***$P < 0.001$.

### Fig. S3

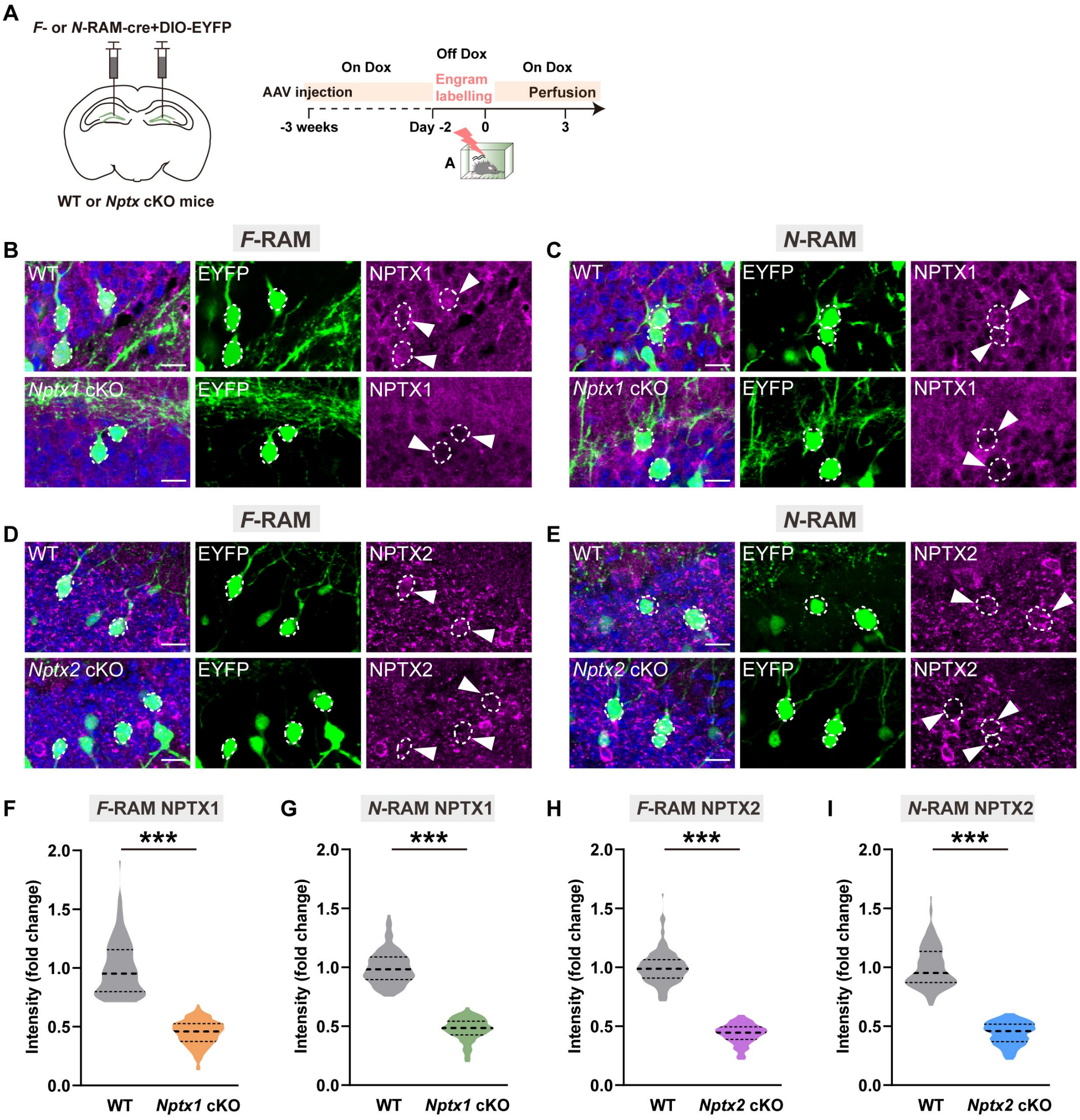
A
F- or N-RAM-cre+DIO-EYFP
WT or Nptx cKO mice
On Dox
Off Dox
On Dox
AAV injection
Engram labelling
Perfusion
-3 weeks
Day -2
0
3
A
B
F-RAM
WT
EYFP
NPTX1
Nptx1 cKO
C
N-RAM
D
NPTX2
Nptx2 cKO
E
F
F-RAM NPTX1
G
N-RAM NPTX1
H
F-RAM NPTX2
I
N-RAM NPTX2
***
Intensity (fold change)
2.0
1.5
1.0
0.5
0.0
WT
Nptx1 cKO
Nptx2 cKO

**Figure S3. IHC validation of NPTX1 and NPTX2 knockout in DG engram ensembles.** (A) Diagram of AAV injection and experimental scheme to label *F*-RAM and *N*-RAM ensembles in WT and *Nptxs* cKO mice. (B, C) Representative confocal images of *F*- or *N*-RAM cells colocalizing with NPTX1. Green: EYFP, Purple: NPTX1, Blue: DAPI. Dashed white lines and white arrows outline cells. Scale bar: 10 μm. (D, E) Representative confocal images of *F*- or *N*-RAM cells colocalizing with NPTX2. Green: EYFP, Purple: NPTX2, Blue: DAPI. Dashed white lines and white arrows outline cells. Scale bar: 10 μm. (F, G) The average NPTX1 fluorescence intensity of *F*- or *N*-RAM cells in WT and *Nptx1* cKO mice. (H, I) The average NPTX2 fluorescence intensity of *F*- or *N*-RAM cells in WT and *Nptx2* cKO mice. Data are presented as mean ± S.E.M; ****P* < 0.001.

Fig. S4

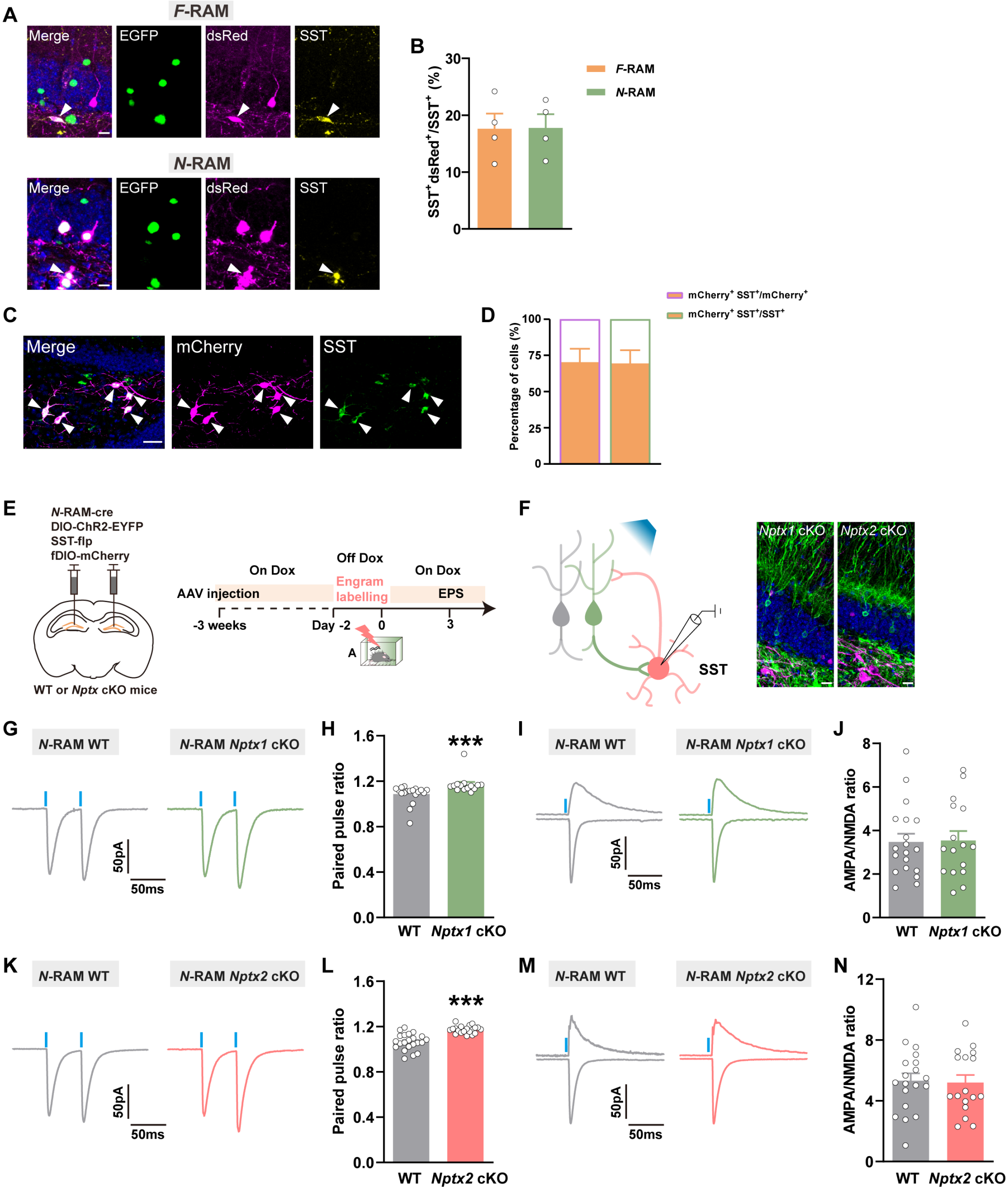

A
F-RAM
Merge
EGFP
dsRed
SST
N-RAM
B
SST+dsRed+/SST+ (%)
F-RAM
N-RAM
C
Merge
mCherry
SST
D
Percentage of cells (%)
mCherry+ SST+/mCherry+
mCherry+ SST+/SST+
E
N-RAM-cre
DIO-ChR2-EYFP
SST-flp
fDIO-mCherry
WT or Nptx cKO mice
AAV injection
On Dox
Off Dox
Engram labelling
On Dox
EPS
-3 weeks
Day -2
0
3
F
SST
Nptx1 cKO
Nptx2 cKO
G
N-RAM WT
N-RAM Nptx1 cKO
50pA
50ms
H
Paired pulse ratio
***
WT
Nptx1 cKO
I
N-RAM WT
N-RAM Nptx1 cKO
J
AMPA/NMDA ratio
WT
Nptx1 cKO
K
N-RAM WT
N-RAM Nptx2 cKO
L
Paired pulse ratio
***
WT
Nptx2 cKO
M
N-RAM WT
N-RAM Nptx2 cKO
N
AMPA/NMDA ratio
WT
Nptx2 cKO

**Figure S4. The effects of *Nptxs* depletion in *N*-RAM ensemble on the plasticity of DG SST$^+$ interneurons.** (A) Representative confocal images of DG local dsRed$^+$ cells immunostaining with SST. Green: EGFP, Purple: dsRed, Yellow: SST, Blue: DAPI. White arrows indicate the dsRed$^+$ SST$^+$ colocalized cells. Scale bar: 5 μm. (B) The percentages of colocalized cells in DG total SST$^+$ cells. (C, D) Representative confocal images and overlap analysis of SST-mCherry colocalizing with SST antibody. Green: SST antibody, Purple: mCherry, Blue: DAPI. Scale bar: 10 μm. N = 6. (E) Diagram of AAV injection and experimental scheme to label *N*-RAM engram ensemble. (F) Diagram of photostimulation and whole-cell patch clamp recordings (left) and representative expression of *N*-RAM engram cells and SST$^+$ interneurons (Right). Green: *N*-RAM engram cells of *Nptx1* and *Nptx2* cKO mice, EYFP, Purple: SST$^+$ interneurons, mCherry, Blue: DAPI. Scale bar: 10 μm. (G, H) Representative traces and quantification of opto-evoked PPR recorded from WT and *Nptx1* cKO mice. (I, J) Representative traces of opto-evoked AMPA-EPSC, NMDA-EPSC and the average A/N ratio recorded from WT and *Nptx1* cKO mice. (K, L) Representative traces and quantification of opto-evoked PPR recorded from WT and *Nptx2* cKO mice. (M, N) Representative traces of opto-evoked AMPA-EPSC, NMDA-EPSC and the average A/N ratio recorded from WT and *Nptx2* cKO mice. Data are presented as mean ± S.E.M; ***$P < 0.001$.

### Fig. S5

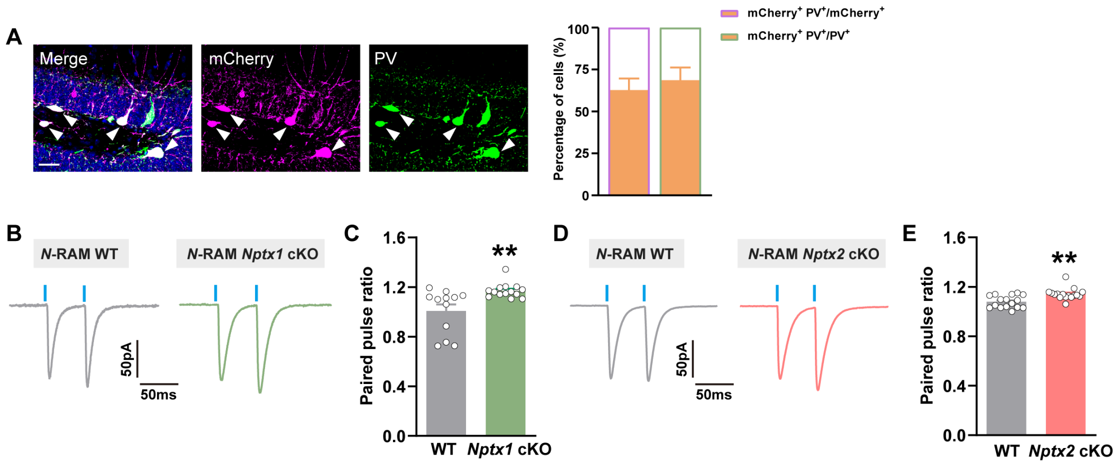
A
Merge
mCherry
PV
Percentage of cells (%)
100
75
50
25
0
mCherry+ PV+/mCherry+
mCherry+ PV+/PV+
B
N-RAM WT
N-RAM Nptx1 cKO
50pA
50ms
C
Paired pulse ratio
1.6
1.2
0.8
0.4
0.0
**
WT
Nptx1 cKO
D
N-RAM WT
N-RAM Nptx2 cKO
50pA
50ms
E
Paired pulse ratio
1.6
1.2
0.8
0.4
0.0
**
WT
Nptx2 cKO

**Figure S5. The effects of *Nptxs* depletion in *N*-RAM ensemble on pre-synaptic glutamate release to DG PV$^+$ interneurons.** (A) Representative confocal images and overlapping analysis of PV-mCherry colocalizing with PV antibody. Green: PV antibody, Purple: mCherry, Blue: DAPI. Scale bar: 10 μm. N = 5. (B, C) Representative traces and quantification of opto-evoked PPR recorded from WT and *Nptx1* cKO mice. (D, E) Representative traces and quantification of opto-evoked PPR recorded from WT and *Nptx2* cKO mice. Data are presented as mean ± S.E.M; $^{**}P < 0.01$.

#### Fig. S6

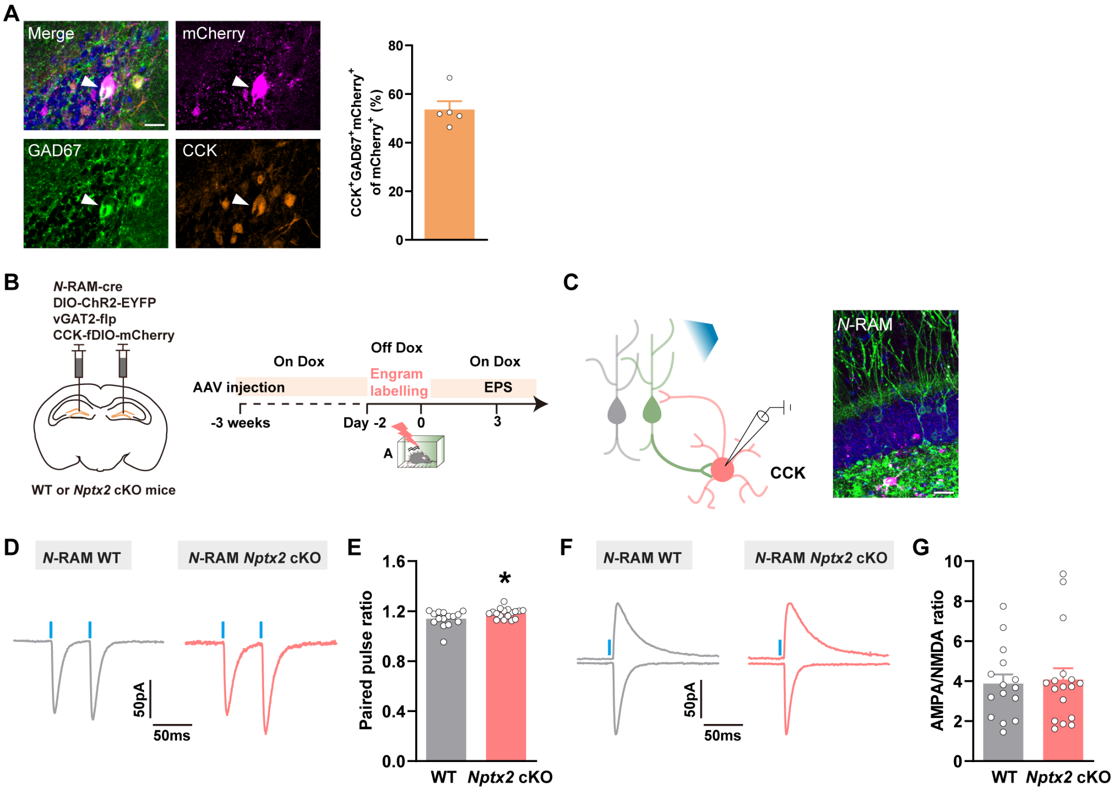

A
Merge
mCherry
GAD67
CCK
CCK+GAD67+mCherry+ of mCherry+ (%)
80
60
40
20
0
B
N-RAM-cre
DIO-ChR2-EYFP
vGAT2-flp
CCK-fDIO-mCherry
WT or Nptx2 cKO mice
On Dox
Off Dox
On Dox
AAV injection
Engram labelling
EPS
-3 weeks
Day -2
0
3
A
C
CCK
N-RAM
D
N-RAM WT
N-RAM Nptx2 cKO
50pA
50ms
E
Paired pulse ratio
1.6
1.2
0.8
0.4
0.0
*
WT
Nptx2 cKO
F
N-RAM WT
N-RAM Nptx2 cKO
50pA
50ms
G
AMPA/NMDA ratio
10
8
6
4
2
0
WT
Nptx2 cKO

**Figure S6. The effects of *Nptxs* depletion in *N*-RAM ensemble on the plasticity of DG CCK$^+$ interneurons.** (A) Representative confocal images and overlap analysis of CCK-mCherry colocalizing with GAD67 and CCK antibody. Green: GAD67 antibody, Yellow, CCK antibody, Purple: mCherry, Blue: DAPI. Scale bar: 10 μm. N = 5. (B) Diagram of AAV injection and experimental scheme to label *N*-RAM engram ensembles. (C) Diagram of photostimulation and whole-cell patch clamp recordings (left) and representative expression of engram cells and CCK$^+$ interneurons (right). Green: *N*-RAM engram cells, EYFP, Purple: CCK$^+$ interneurons, mCherry, Blue: DAPI. Scale bar: 10 μm. (D, E) Representative traces and quantification of opto-evoked PPR recorded from WT and *Nptx2* cKO mice. (F, G) Representative traces of opto-evoked AMPA-EPSC, NMDA-EPSC and the average A/N ratio recorded from WT and *Nptx2* cKO mice. Data are presented as mean ± S.E.M; $*P < 0.05$.

### Fig. S7

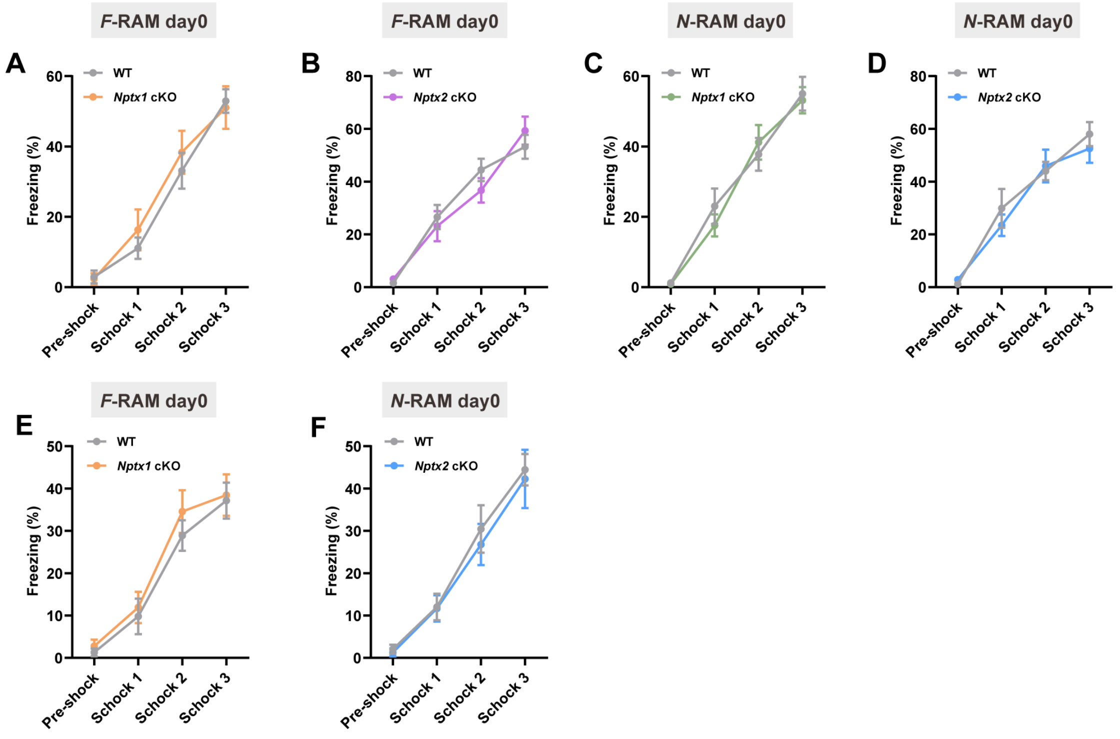

F-RAM day0
N-RAM day0
A
B
C
D
E
F
WT
Nptx1 cKO
Nptx2 cKO
Freezing (%)
Pre-shock
Schock 1
Schock 2
Schock 3

**Figure S7. Freezing level during fear conditioning and cell counts of *F*-RAM and *N*-RAM engram cells.** (A) The quantification for freezing levels of WT and *Nptx1* cKO mice during CFC (*F*-RAM). (B) The quantification for freezing levels of WT and *Nptx2* cKO mice during CFC (*F*-RAM). (C) The quantification for freezing levels of WT and *Nptx1* cKO mice during CFC (*N*-RAM). (D) The quantification for freezing levels of WT and *Nptx2* cKO mice during CFC (*N*-RAM). (E) The quantification for freezing levels of WT and *Nptx1* cKO mice during CFC (*F*-RAM). (F) The quantification for freezing levels of WT and *Nptx2* cKO mice during CFC (*N*-RAM). Data are presented as mean ± S.E.M.

#### Fig. S8

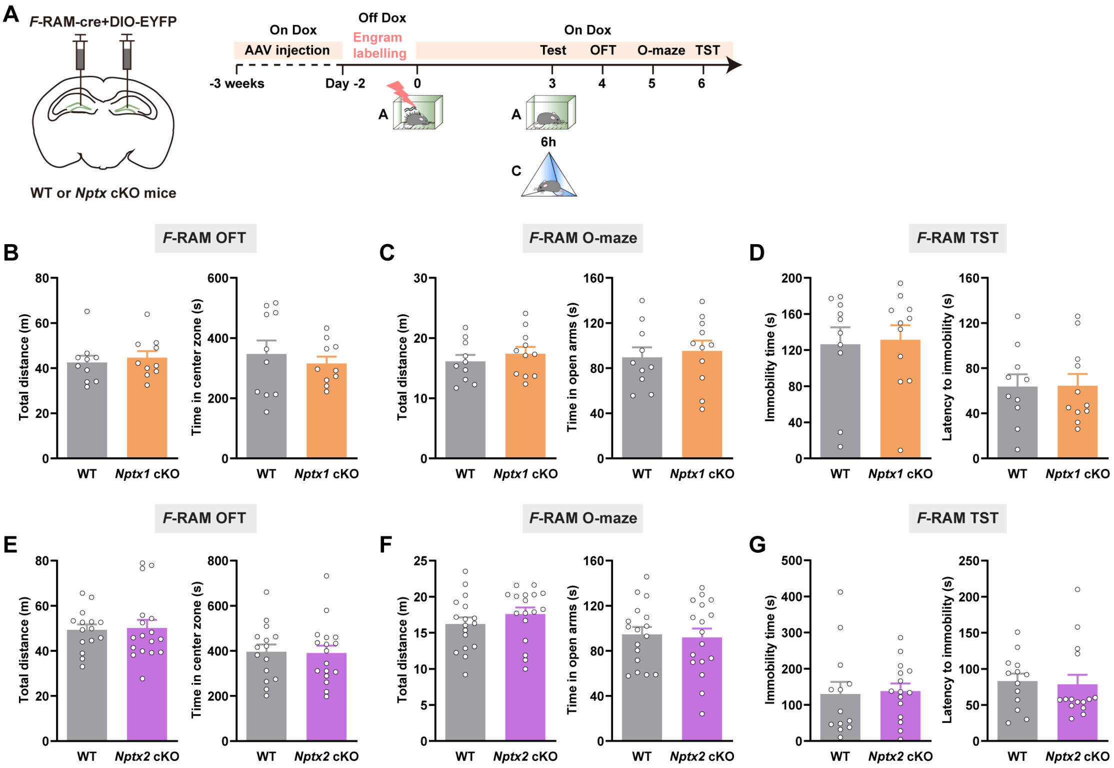

A
F-RAM-cre+DIO-EYFP
WT or Nptx cKO mice
On Dox
AAV injection
Off Dox
Engram labelling
On Dox
Test
OFT
O-maze
TST
-3 weeks
Day -2
0
3
4
5
6
A
A
6h
C
F-RAM OFT
F-RAM O-maze
F-RAM TST
B
C
D
E
F
G
Total distance (m)
Time in center zone (s)
Time in open arms (s)
Immobility time (s)
Latency to immobility (s)
WT
Nptx1 cKO
Nptx2 cKO

**Figure S8. The effects of *Nptxs* depletion in *F*-RAM ensemble activated by CFC on the locomotion, anxiety or depression level of mice.** (A) Diagram of AAV injection and experimental scheme of open field test (OFT), O-maze test and tail suspension test (TST). (B) The average distance travelled (left) and time spent in center zone (right) of OFT for WT and *Nptx1* cKO mice. (C) The average distance travelled (left) and time spent in open arms (right) of O-maze test for WT and *Nptx1* cKO mice. (D) The average immobility time (left) and latency to immobility (right) of TST for WT and *Nptx1* cKO mice. (E) The average distance travelled (left) and time spent in center zone (right) of OFT for WT and *Nptx2* cKO mice. (F) The average distance travelled (left) and time spent in open arms (right) of O-maze test for WT and *Nptx2* cKO mice. (G) The average immobility time (left) and latency to immobility (right) of TST for WT and *Nptx2* cKO mice. Data are presented as mean ± S.E.M.

### Fig. S9

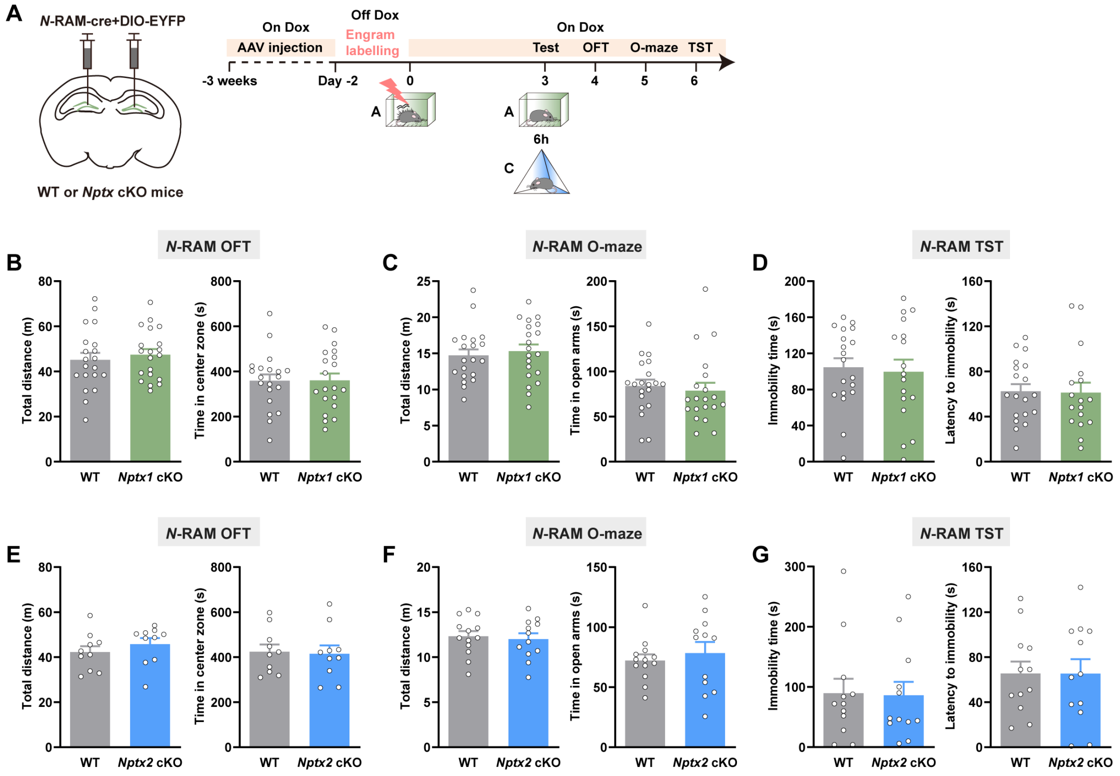

A
N-RAM-cre+DIO-EYFP
WT or Nptx cKO mice
On Dox
AAV injection
Off Dox
Engram labelling
On Dox
Test
OFT
O-maze
TST
-3 weeks
Day -2
0
3
4
5
6
A
A
6h
C
B
N-RAM OFT
Total distance (m)
Time in center zone (s)
WT
Nptx1 cKO
C
N-RAM O-maze
Total distance (m)
Time in open arms (s)
D
N-RAM TST
Immobility time (s)
Latency to immobility (s)
E
N-RAM OFT
Nptx2 cKO
F
N-RAM O-maze
G
N-RAM TST

**Figure S9. The effects of Nptxs depletion in N-RAM ensemble activated by CFC on the locomotion, anxiety or depression level of mice.** (A) Diagram of AAV injection and experimental scheme of OFT, O-maze test and TST. (B) The average distance travelled (left) and time spent in center zone (right) of OFT for WT and Nptx1 cKO mice. (C) The average distance travelled (left) and time spent in open arms (right) of O-maze test for WT and Nptx1 cKO mice. (D) The average immobility time (left) and latency to immobility (right) of TST for WT and Nptx1 cKO mice. (E) The average distance travelled (left) and time spent in center zone (right) of OFT for WT and Nptx2 cKO mice. (F) The average distance travelled (left) and time spent in open arms (right) of O-maze test for WT and Nptx2 cKO mice. (G) The average immobility time (left) and latency to immobility (right) of TST for WT and Nptx2 cKO mice. Data are presented as mean ± S.E.M.

### Fig. S10

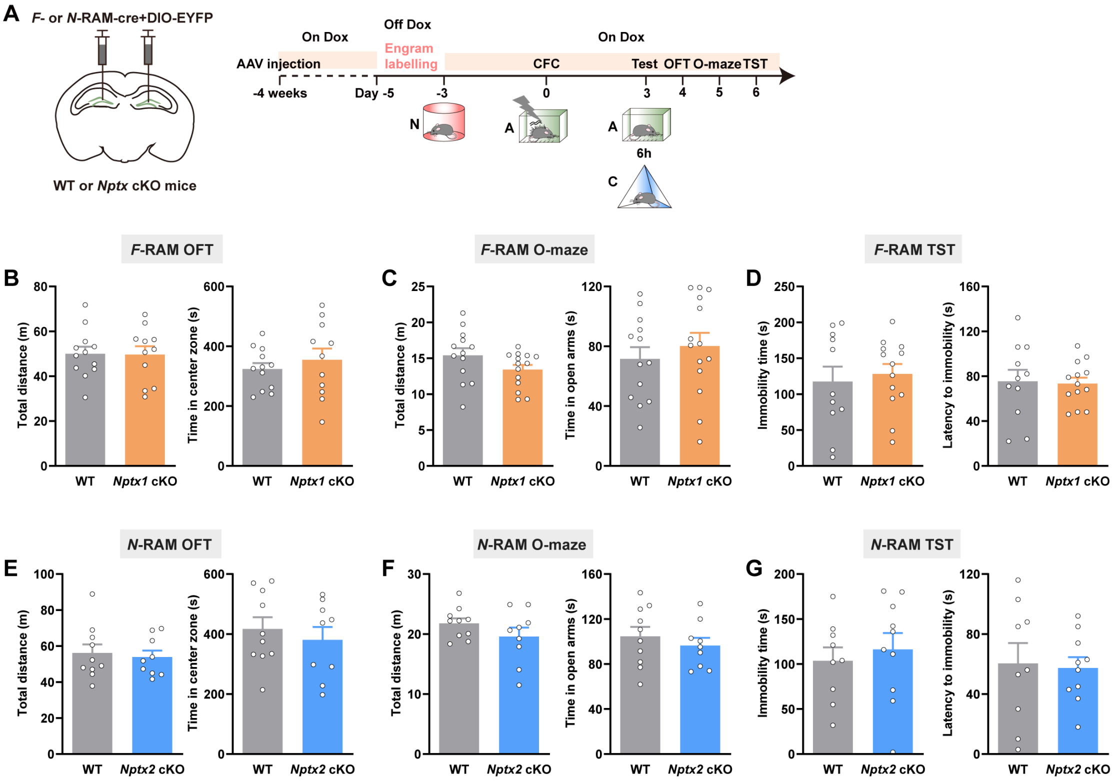
A
F- or N-RAM-cre+DIO-EYFP
WT or Nptx cKO mice
On Dox
Off Dox
On Dox
AAV injection
Engram labelling
CFC
Test OFT O-maze TST
-4 weeks
Day -5
-3
0
3
4
5
6
N
A
A
6h
C
F-RAM OFT
F-RAM O-maze
F-RAM TST
B
C
D
Total distance (m)
Time in center zone (s)
Time in open arms (s)
Immobility time (s)
Latency to immobility (s)
WT
Nptx1 cKO
N-RAM OFT
N-RAM O-maze
N-RAM TST
E
F
G
Nptx2 cKO

**Figure S10. The effects of *Nptxs* depletion in *F*-and *N*-RAM ensembles activated by novel context exposure on the locomotion, anxiety or depression level of mice.** (A) Diagram of AAV injection and experimental scheme of OFT, O-maze test and TST. (B) The average distance travelled (left) and time spent in center zone (right) of OFT for WT and *Nptx1* cKO mice (*F*-RAM). (C) The average distance travelled (left) and time spent in open arms (right) of O-maze test for WT and *Nptx1* cKO mice (*F*-RAM). (D) The average immobility time (left) and latency to immobility (right) of TST for WT and *Nptx1* cKO mice (*F*-RAM). (E) The average distance travelled (left) and time spent in center zone (right) of OFT for WT and *Nptx2* cKO mice (*N*-RAM). (F) The average distance travelled (left) and time spent in open arms (right) of O-maze test for WT and *Nptx2* cKO mice (*N*-RAM). (G) The average immobility time (left) and latency to immobility (right) of TST for WT and *Nptx2* cKO mice (*N*-RAM). Data are presented as mean ± S.E.M.

Fig. S11

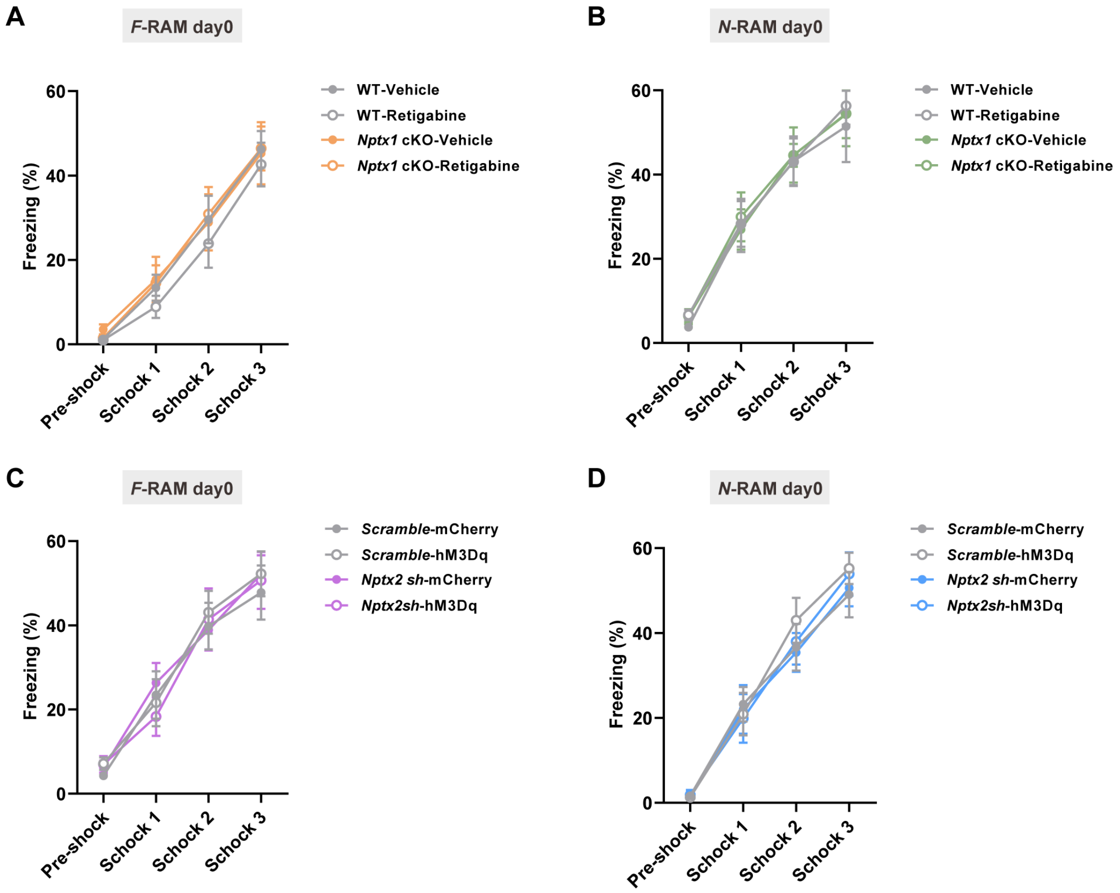
A
F-RAM day0
Freezing (%)
0
20
40
60
Pre-shock
Schock 1
Schock 2
Schock 3
WT-Vehicle
WT-Retigabine
Nptx1 cKO-Vehicle
Nptx1 cKO-Retigabine
B
N-RAM day0
Freezing (%)
0
20
40
60
Pre-shock
Schock 1
Schock 2
Schock 3
WT-Vehicle
WT-Retigabine
Nptx1 cKO-Vehicle
Nptx1 cKO-Retigabine
C
F-RAM day0
Freezing (%)
0
20
40
60
Pre-shock
Schock 1
Schock 2
Schock 3
Scramble-mCherry
Scramble-hM3Dq
Nptx2 sh-mCherry
Nptx2sh-hM3Dq
D
N-RAM day0
Freezing (%)
0
20
40
60
Pre-shock
Schock 1
Schock 2
Schock 3
Scramble-mCherry
Scramble-hM3Dq
Nptx2 sh-mCherry
Nptx2sh-hM3Dq

**Figure S11. The effects of activating Kv7.2 or $PV^+$ interneurons in DG on the freezing levels of mice during CFC.** (A, B) The quantitative analysis for freezing levels of WT-Vehicle, WT-Retigabine, *Nptx1* cKO-Vehicle and *Nptx1* cKO-Retigabine mice during CFC. (C, D) The quantitative analysis for freezing levels in *Scramble*-mCherry, *Scramble*-hM3Dq, *Nptx2 sh*-mCherry and *Nptx2 sh*-hM3Dq groups during CFC. Data are presented as mean ± S.E.M.

#### Fig. S12

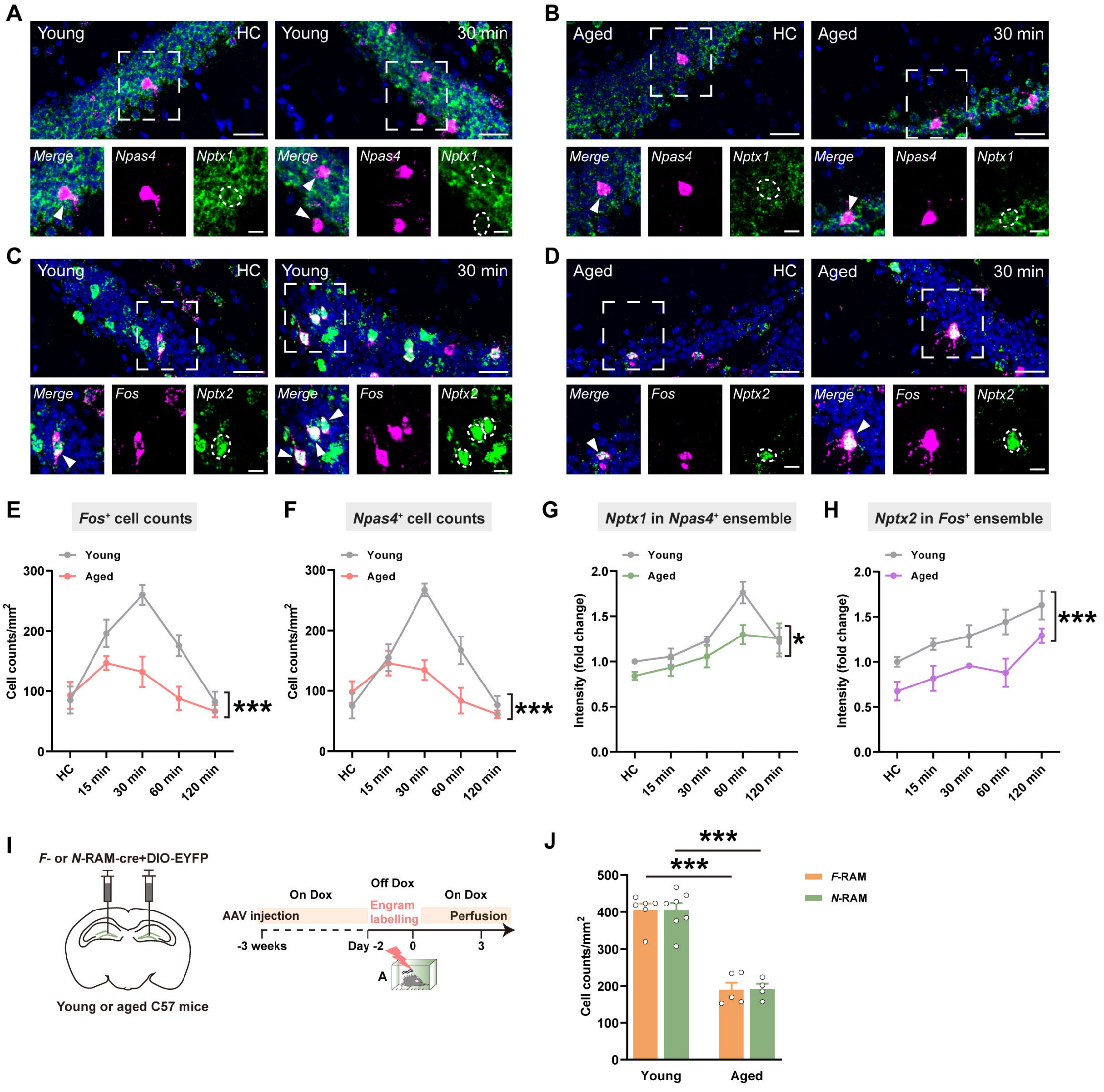

A
Young
HC
30 min
Merge
Npas4
Nptx1
B
Aged
C
Fos
Nptx2
D
E
Fos+ cell counts
F
Npas4+ cell counts
G
Nptx1 in Npas4+ ensemble
H
Nptx2 in Fos+ ensemble
Cell counts/mm2
Intensity (fold change)
15 min
60 min
120 min
***
*
I
F- or N-RAM-cre+DIO-EYFP
Young or aged C57 mice
On Dox
Off Dox
AAV injection
Engram labelling
Perfusion
-3 weeks
Day -2
0
3
J
F-RAM
N-RAM

**Figure S12. Engram cell counts and *Nptxs* expression in DG *F*-RAM and *N*-RAM ensembles in young and aged mice.** (A, B) Representative confocal images of *Nptx1* colocalizing with *Npas4* under HC condition and 30 min after CFC in DG of young and aged mice. Green: *Nptx1*, Purple: *Npas4*, Blue: DAPI. White arrows indicate the colocalized cells. Scale bar: top, 30 μm, bottom, 10 μm. (C, D) Representative confocal images of *Nptx2* colocalizing with *Fos* under HC condition and 30 min after CFC in DG of young and aged mice. Green: *Nptx2*, Purple: *Fos*, Blue: DAPI. White arrows indicate the colocalized cells. Scale bar: top, 30 μm, bottom, 10 μm. (E, F) The *Fos*$^{+}$ and *Npas4*$^{+}$ cell counts in DG. (G) The fluorescence intensity of *Nptx1* mRNA in *Npas4*$^{+}$ ensemble at HC, 15 min, 30 min, 60 min and 120 min after CFC. (H) The fluorescence intensity of *Nptx2* mRNA in *Fos*$^{+}$ ensemble at HC, 15 min, 30 min, 60 min and 120 min after CFC. (I) Diagram of AAV injection and experimental scheme to label *F*-RAM and *N*-RAM cells, (J) The *F*-RAM and *N*-RAM cell counts in DG of young and aged mice. Data are presented as mean ± S.E.M; *$P < 0.05$, ***$P < 0.001$.

#### Fig. S13

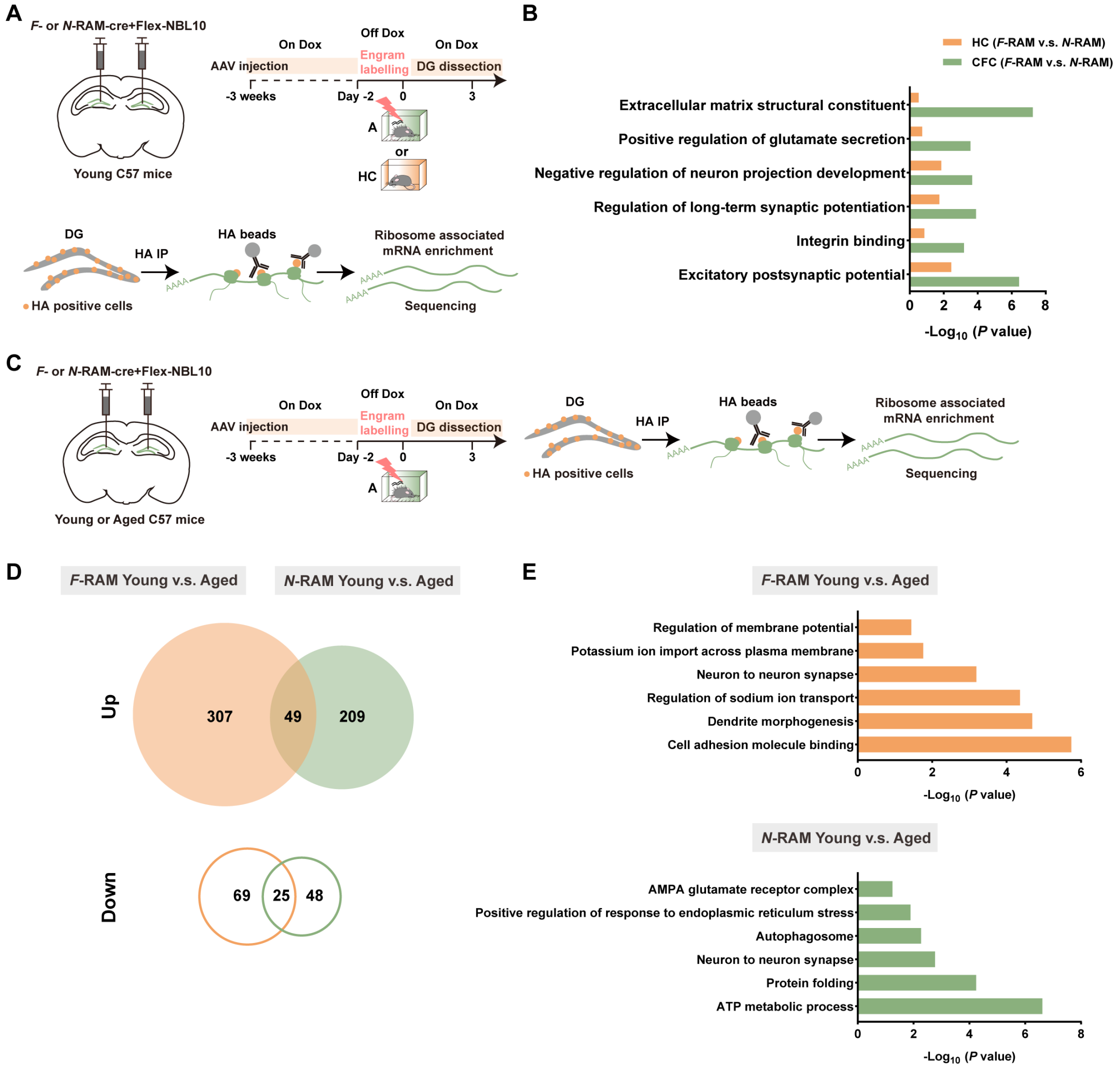
A
F- or N-RAM-cre+Flex-NBL10
Young C57 mice
On Dox
Off Dox
On Dox
AAV injection
Engram labelling
DG dissection
-3 weeks
Day -2
0
3
A
or
HC
DG
HA IP
HA beads
Ribosome associated mRNA enrichment
HA positive cells
Sequencing
B
HC (F-RAM v.s. N-RAM)
CFC (F-RAM v.s. N-RAM)
Extracellular matrix structural constituent
Positive regulation of glutamate secretion
Negative regulation of neuron projection development
Regulation of long-term synaptic potentiation
Integrin binding
Excitatory postsynaptic potential
-Log10 (P value)
C
F- or N-RAM-cre+Flex-NBL10
Young or Aged C57 mice
D
F-RAM Young v.s. Aged
N-RAM Young v.s. Aged
Up
307
49
209
Down
69
25
48
E
F-RAM Young v.s. Aged
Regulation of membrane potential
Potassium ion import across plasma membrane
Neuron to neuron synapse
Regulation of sodium ion transport
Dendrite morphogenesis
Cell adhesion molecule binding
N-RAM Young v.s. Aged
AMPA glutamate receptor complex
Positive regulation of response to endoplasmic reticulum stress
Autophagosome
Neuron to neuron synapse
Protein folding
ATP metabolic process
-Log10 (P value)

**Figure S13. Transcriptional networks enrichment in *F*-RAM and *N*-RAM ensembles in young and aged mice.** (A) Diagram of AAV injection, experimental scheme to label *F*-RAM and *N*-RAM ensembles and scheme of RiboTag enrichment of *F*- and *N*-RAM transcriptomes activated in HC or by CFC of young mice. (B) GO enrichment analysis of transcriptional differences in DG *F*-RAM and *N*-RAM ensembles activated in HC or by CFC of young mice. *F*-RAM, HC, n = 4. *N*-RAM, HC, n = 4. *F*-RAM, CFC, n = 4. *N*-RAM, CFC, n = 4. (C) Diagram of AAV injection, experimental scheme to label *F*-RAM and *N*-RAM ensembles and scheme of RiboTag enrichment of *F*- and *N*-RAM transcriptomes in young and aged mice. (D) A Venn diagram showing the upregulation (up) and downregulation (bottom) of *F*- and *N*-RAM transcriptomes during aging and the overlap between these two groups. (E) GO enrichment analysis of transcriptional differences in DG *F*-RAM (up) and *N*-RAM (down) ensembles respectively along with aging. *F*-RAM, young, n = 5. *F*-RAM, aged, n = 5. *N*-RAM, young, n = 6. *N*-RAM, aged, n = 7. Data are presented as mean ± S.E.M.

### Fig. S14

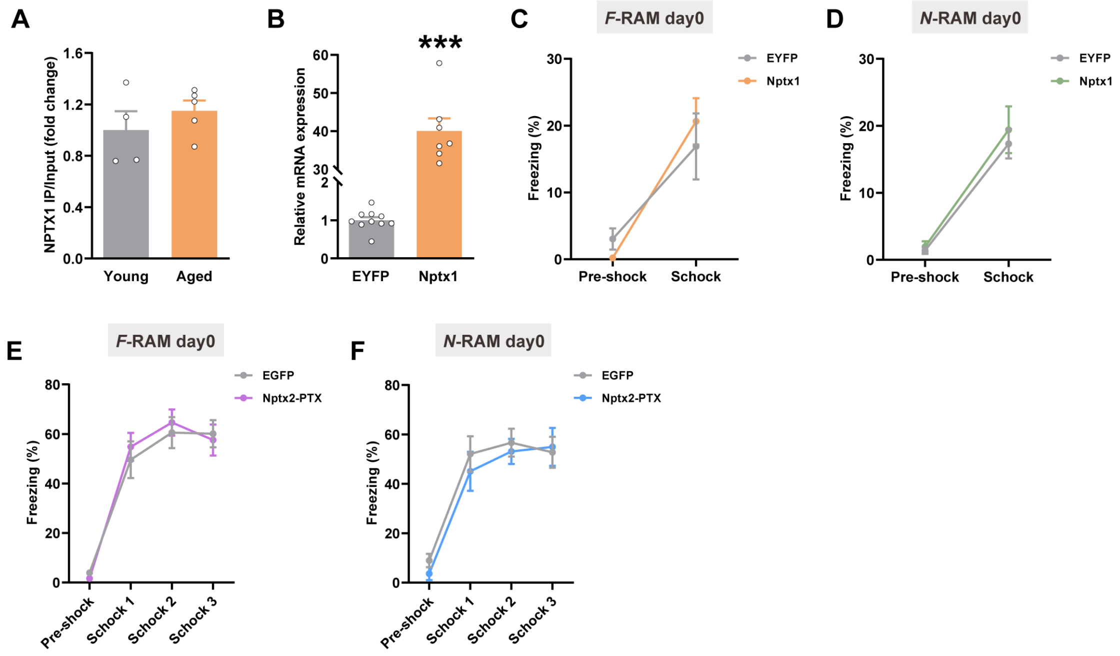
A
NPTX1 IP/Input (fold change)
1.6
1.2
0.8
0.4
0.0
Young
Aged
B
Relative mRNA expression
60
50
40
30
2
1
0
***
EYFP
Nptx1
C
F-RAM day0
Freezing (%)
30
20
10
0
EYFP
Nptx1
Pre-shock
Schock
D
N-RAM day0
Freezing (%)
30
20
10
0
EYFP
Nptx1
Pre-shock
Schock
E
F-RAM day0
Freezing (%)
80
60
40
20
0
EGFP
Nptx2-PTX
Pre-shock
Schock 1
Schock 2
Schock 3
F
N-RAM day0
Freezing (%)
80
60
40
20
0
EGFP
Nptx2-PTX
Pre-shock
Schock 1
Schock 2
Schock 3

**Figure S14. The effects of *Nptxs* overexpression in aged mice on the freezing level during conditioning.** (A) The quantification of IP-NPTX1 in young and aged mice. (B) RT-qPCR analysis of Nptx1 overexpression efficiency. (C, D) The average freezing levels of EYFP and Nptx1 aged mice during CFC. (E, F) The average freezing levels of EYFP and Nptx2-PTX aged mice during CFC. Data are presented as mean ± S.E.M; ***$P$ < 0.001.

### Fig. S15

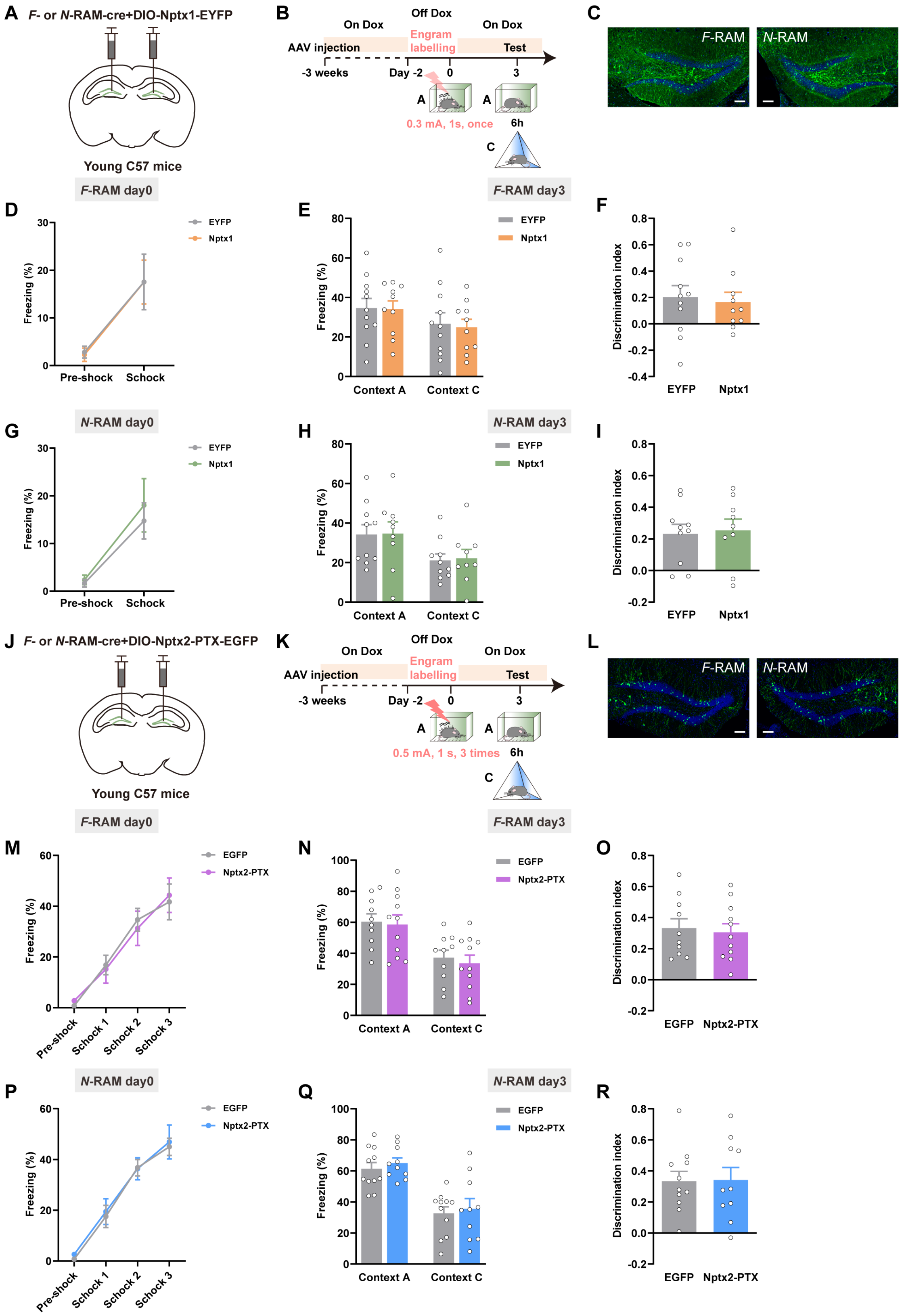
A F- or N-RAM-cre+DIO-Nptx1-EYFP
Young C57 mice
B On Dox
Off Dox
On Dox
AAV injection
Engram labelling
Test
-3 weeks
Day -2
0
3
0.3 mA, 1s, once
6h
C F-RAM
N-RAM
F-RAM day0
F-RAM day3
D Freezing (%)
Pre-shock
Schock
EYFP
Nptx1
E Freezing (%)
Context A
Context C
F Discrimination index
N-RAM day0
N-RAM day3
G
H
I
J F- or N-RAM-cre+DIO-Nptx2-PTX-EGFP
K 0.5 mA, 1 s, 3 times
L
M Pre-shock
Schock 1
Schock 2
Schock 3
EGFP
Nptx2-PTX
N
O
P
Q
R

**Figure S15. The effects of overexpressing NPTXs in DG engram ensembles on memory expression in young mice.** (A, J) Diagram of AAV injection. (B, K) Experimental scheme of memory retrieval test. (C, L) Representative expression of NPTX1 or NPTX2-PTX in *F*- or *N*-RAM engram cells in DG. Green: *F*-RAM or *N*-RAM ensemble, EYFP or EGFP, Blue: DAPI. Scale bar: 100 μm. (D, G) The quantitative analysis for freezing levels of EYFP and Nptx1 young mice during CFC. (E, F) The freezing percentage and discrimination index of EYFP and Nptx1 young groups (*F*-RAM). (H, I) The freezing percentage and discrimination index of EYFP and Nptx1 young groups (*N*-RAM). (M, P) The quantitative analysis for freezing levels of EGFP and NPTX2-PTX young mice during CFC. (N, O) The freezing percentage and discrimination index of EGFP and Nptx2-PTX young groups (*F*-RAM). (Q, R) The freezing percentage and discrimination index of EGFP and Nptx2-PTX young groups (*N*-RAM). Data are presented as mean ± S.E.M.
